# Connectome of a model local cortical circuit flexibly shapes layer-dependent multi-frequency oscillations

**DOI:** 10.1101/026674

**Authors:** Markus Helmer, Xue Jie Chen, Wei Wei, Fred Wolf, Demian Battaglia

## Abstract

The role played by interlayer connections in shaping local responses and their long-range coupling has not yet been fully elucidated. Here, we analyze a rate model of a canonic local circuit with realistic anatomy. We find that this circuit generates a rich repertoire of possible dynamical states, including an oscillatory regime in which gamma-and beta-oscillations dominate in superficial and deep layers, respectively, in agreement with experimental observations. This regime stems from non-linear inter-layer interactions, independently from intrinsic resonance properties of distinct layers. Moreover, by connecting two local circuits via cortico-cortical projections, the emergent phase differences define a flexible and frequency-dependent inter-areal hierarchy. Such dynamic patterns generally do not arise in randomized circuits, and the compatible connectomes are rare, although not unique. Altogether, these results suggest that inter-layer connectivity is homeostatically regulated to make local circuits fit to integrate and multiplex signals from several sources in multiple frequency bands.

**Author Summary:** The local circuit of mammalian cortex presents a characteristic multilayered structure, with feedforward (and feedback) cortico-cortical connections originating from (and targeting) distinct and well defined layers. Here, we model how such a structure fundamentally shapes the dynamical repertoire of local cortical oscillatory states and their long-range interaction. Experimental evidence, matched by our simulations, suggests that different cortical layers oscillate at different frequencies and that neuronal oscillations at different frequencies are exploited for communication in different directions. While this laminar specificity of oscillations is often explained in terms of multiple inhibitory populations with different resonance properties, we show here that it could alternatively emerge as a byproduct of the collective local circuit dynamics. Our modelling study indicates furthermore that the empirically observed multi-frequency oscillatory patterns cannot be reproduced in presence of an arbitrary interlayer connectivity. In this sense, therefore, we believe that the adopted connectome, derived from neuroanatomical reconstructions, is “special”. Nevertheless, it is not unique, since other, very different connectomes may also lead to a matching dynamical repertoire. This suggests that a multiplicity of non-random canonical circuit templates may share largely overlapping functions, robustly achieved and maintained via functional homeostasis mechanisms.

## Introduction

The microcircuits of mammalian brain cortex present prominent hallmark features such as a six-layered architecture [1] and an organization into vertical columns [2]. Remarkable regularities in the wiring between neuronal populations in different layers, both within and between columns [3–8] have led to propose the existence of a “canonical local microcircuit”, providing a building block for larger scale cortical networks. This hypothesis culminated in the compilation of quantitative interlayer wiring diagrams [9–11], which despite their approximations and limits [12], have proven particularly attractive, bolstering theories about specialized roles of different layers [13–15] as well as motivating computational investigations of the local microcircuit function [11, 16–20].

The layered structure of cortical circuits also affects the generation and the properties of brain oscillations. On the one hand, ascending and descending structural connections originate from more superficial and deep layers, respectively [7, 21]. On the other hand, oscillations of neural activity in different layers have different spatial spread and power [22, 23] and display spectral resonances at layer-dependent frequencies [24–27], with gamma band oscillations arising in more superficial layers and slower oscillations dominant in deeper layers [24, 26]. Correspondingly, it was proposed that inter-areal communication in different directions exploits oscillatory coherence in different frequency bands [28–32]: gamma-band frequencies for bottom-up interactions, and alpha-/beta-band frequencies for top-down interactions, matching the predominant oscillations in the connections’ source layers.

However, these correlational observations do not allow to determine whether local microcircuit connectivity plays an actual causal role in shaping layer-specific properties of neural oscillations and of their flexible modulation by context. To demonstrate the existence of a direct influence we adopt here a computational approach. Through systematic simulations of a rate model embedding an anatomically realistic multi-layer connectivity [10], we reveal that interlayer interactions give rise to a rich repertoire of possible oscillatory modes. This dynamical repertoire includes robust regimes in which the laminar separation between slower and faster oscillations emerges purely through network mechanisms, not necessarily requiring the introduction of neuronal populations with distinct intrinsic resonance frequencies, unlike in other models [33].

Furthermore, we couple together multiple local circuits, mimicking the laminar arrangement of feedforward and feedback long-range projections established between regions at different levels in the hierarchy of cortical areas [21, 34, 35]. We observe that the dynamics of the interacting local circuits, without need of further ad hoc mechanisms, lead to the self-organized emergence of out-of-phase locking relations between the locally generated multi-frequency rhythms, which are compatible with the already mentioned frequency-specificity of top-down and bottom-up functional interactions [31, 32].

Finally, we inquire whether the observed effect of interlayer connectivity on multi-frequency oscillatory activity is to be attributed to unique properties of the adopted connectome, or whether it is a general outcome of unspecific interlayer interactions. To do so, we compare the self-organized oscillatory dynamics of our model with realistic anatomy with the one generated by models with randomized connectomes. We find that random connectomes do not lead to a natural layer separation between slower and fast oscillations. On the contrary, we are able to generate such a dynamical regime only by carefully selecting the interlayer wiring among a highly restricted set of connectivity configurations, which include the reference connectome by [10], but other very diverse connectomes as well. Therefore, despite this lack of uniqueness, we provide evidence that the empirical canonical microcircuit belongs at least to a very exclusive club of degenerate structures all achieving a common target behavior, in a way reminiscent of variability compensation and homeostasis mechanisms in other neuronal systems [36–38]. This suggests the intriguing possibility that specific micro-circuit patterns are selectively targeted and maintained through evolution and development, because of the functional advantages for inter-areal communication [39] that the associated multi-frequency oscillatory “dynome” [40] confers.

## Results

### Rate model of the canonical local circuit

We analyzed the dynamics of a rate model of a local cortical circuit, whose realistic connectivity, illustrated in Figure 1, was inspired from anatomical studies. It consisted of five layers (L1, L2/3, L4, L5 and L6), each containing one excitatory and one inhibitory population-unit which represented the mean activity in the corresponding population. The connection weights between all populations were taken to be proportional to the relative number of synapses between excitatory (E) and inhibitory (I) populations as measured by [10] (see Figure 1A–D). The actual strengths of inter-population connections were then obtained by multiplying these relative numbers by two phenomenological parameters, *K*_E_ and *K*_I_, indicating global scales of the strengths of excitatory and inhibitory connections, respectively. We assumed here, that these parameters depended only on the type of connection (E or I) but were otherwise independent of the target population. Note that ref. [10] did not report outgoing connections from L1. Therefore, we ignored L1 in the following, given its lack of influence on the dynamics of the other layers, reducing correspondingly the number of neuronal populations explicitly included in the model to eight.

**Figure 1.**
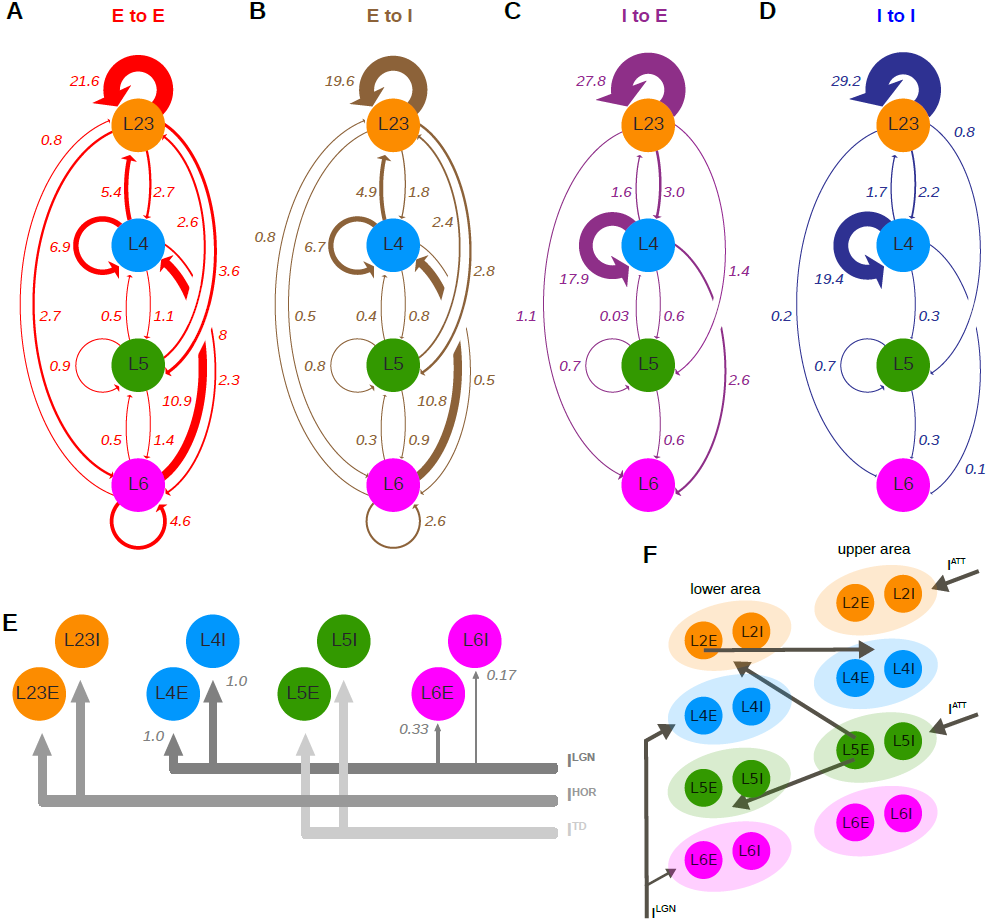
Rate model for a local cortical circuit with realistic inter-layer connectivity. We simulated a canonic cortical local circuit, considering one excitatory (E) and one inhibitory (I) population per layer and connecting them via delayed interactions using experimentally determined relative weights from [10] (reproduced along the arrows in **A-D**). **(E)** Contextual bottom-up (“LGN”), horizontal (“HOR”) and top-down (“TD”) influences extrinsic to the local circuit targeted specific populations in accordance with their known layer-specificity (LGN input was weighted following [10]). **(F)** We also analyzed the interaction of two local circuits assumed to reside at different stages of the cortical hierarchy. Connections between areas had a longer delay than within a column, and an ad hoc determined strength. Contextual inputs (“ATT”) targeted the upper column to emulate activation of a top-down modulating brain state (e.g. attention).

Connections in our model did not give rise to instantaneous interactions, but were delayed by a time *D*, assumed for simplicity to be identical for intra- and interlayer monosynaptic connections. The inclusion of delays favored the emergence of oscillations in the firing rates when the column was stimulated by a constant baseline background current *I*^BG^ (received by all populations). Additional external inputs were targeting specific layers (Figure 1E). As the anatomical connections were measured in area 17, bottom-up input *I*^LGN^ from LGN was sent to L4 and, to a lesser extent, L6, consistent with the literature [7, 10, 34, 41]. Likewise, we mimicked horizontal connections from other columns within the same cortical region via an input *I*^HOR^ specific to L2/3 and “top-down” connections from extra-striate cortices via an input *I*^TD^ specific to L5. By varying the levels of these inputs and their relative balance, different perceptual and cognitive contexts can be phenomenologically emulated. For instance, increasing *I*^LGN^ can represent an increase of contrast of a centrally presented stimulus, and enhanced *I*^HOR^ or *I*^TD^ the presence of modulatory signals from, respectively, the classical or the extra-classical surround of the local circuit receptive field [42]. Alternatively, top-down modulatory inputs from higher order cortical areas, such as, e.g., an “attentional spotlight” sent by prefrontal areas [43] may be represented by a simultaneous increase of both *I*^TD^ and *I*^HOR^ (analogously to [18]).

### Model generates a rich dynamical repertoire

For each fixed value of the background and external inputs, and depending on the efficacies of excitatory and inhibitory interconnections, the model local circuit gave rise to a large repertoire of different possible dynamical states, including steady, oscillatory or chaotic firing modes. To assess the behavior for different values of *K*_E_ and *K*_I_, we simulated the activity of the model in different regions of the parameter space and systematically extracted four summary statistics (see Materials and Methods) for each layer which, taken together, provided a qualitative profile of the dynamical regime. The first metric was the mean firing rate. The second was the relative fraction of low power, defined as a layer’s integrated power spectrum below 30 Hz, *P*_lo_, divided by the summed power spectrum over all frequencies, *P*_tot_. The amplitude and the lag of the peaks (excluding the central, zero-delay one) of the autocorrelation of each layer’s trace provided then the other two metrics. The lag of the first peak measured the period of the fastest appreciable oscillatory structure in the time trace (for easier assessment we present it as a multiple of L4E’s delay), whereas the value of the highest peak in the autocorrelation quantified the degree to which the trace was either more periodic (autocorrelation has values close to 1) or more chaotic (autocorrelation values close to 0). Plots of these quantities in dependence of *K*_E_ and *K*_I_, which we will refer to as *dynamic regime profiles*, summarize the behavior of the model circuit.

We first studied the behavior of the column in a condition of exclusive bottom-up drive (*I*^LGN^ = 2, *I*^HOR^ = *I*^TD^ = 0), besides the always present background input. The dynamic regime profiles of Figure 2A (and see Figs. S1 and S2 for other input configurations) show that while firing rates varied smoothly with *K*_E_ and *K*_I_, sharp transitions were visible in the three other monitored metrics, revealing the existence of homogeneous regimes, i.e. regions of the parameter space with qualitatively distinct but internally uniform dynamics.

**Figure 2.**
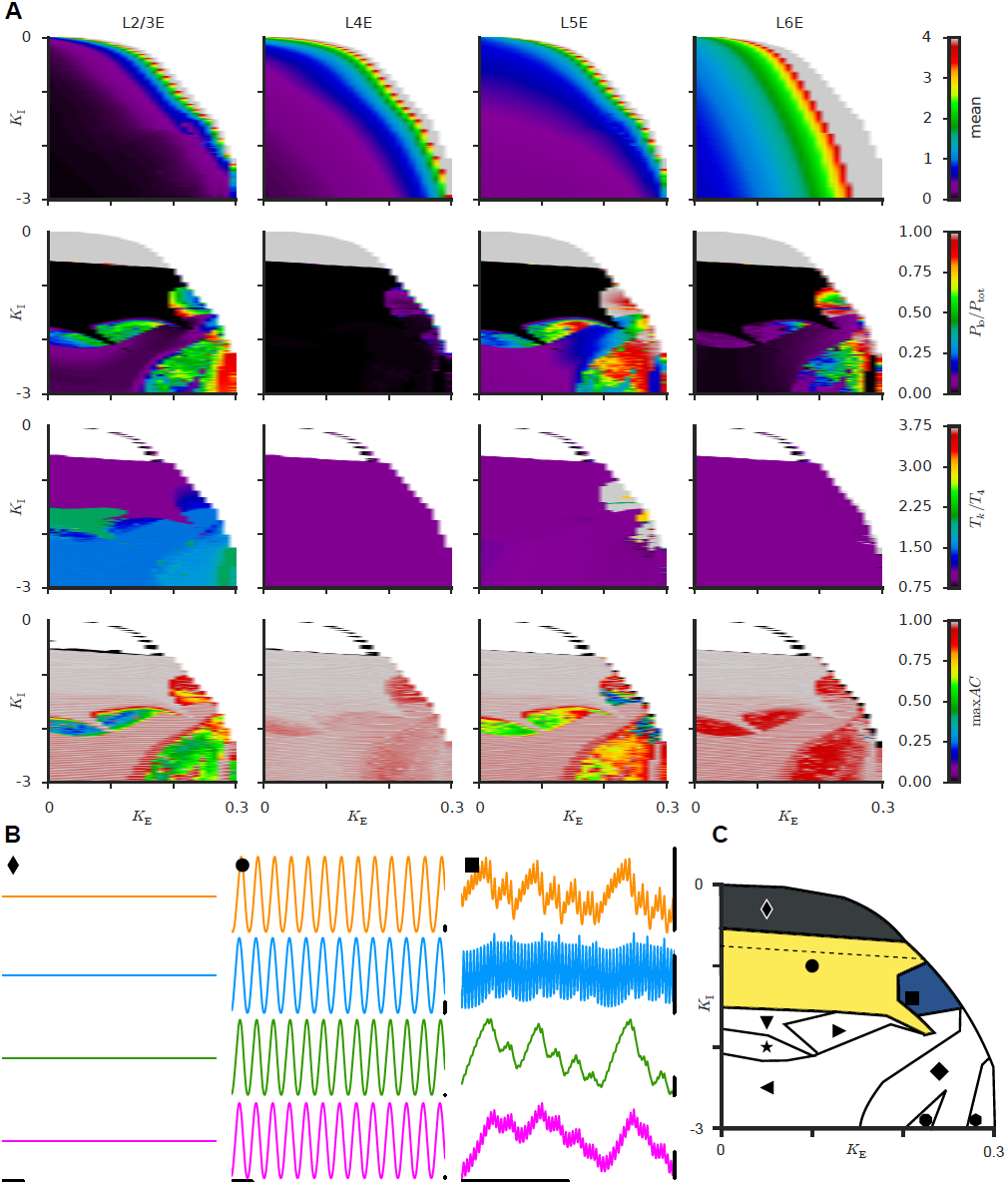
*(following page)* Model yields various qualitatively different possible dynamics. The model behavior was studied in dependence on two phenomenological parameters, *K*_E_ and *K*_I_, multiplying relative excitatory and inhibitory weights provided by Fig. 1, respectively, to set the absolute connection strength. **(A)** Parameter-dependence profiles for different summary statistics. Columns refer to different layers. Rows show from top to bottom: mean activity; relative fraction of low power (signaling, in oscillatory regimes, presence of slow oscillatory components); time lag of autocorrelation’s first peak, in units of layer 4’s (signaling, in oscillatory regimes, the period of main oscillatory component, relative to L4 fast rhythm); and the amplitude of autocorrelation’s highest non-zero-lag peak (with small amplitudes signaling temporally irregular or chaotic waveforms). Such dynamic profiles reveal qualitatively different dynamics depending on efficacy of excitation (*K*_E_) and inhibition (*K*_I_), with inferred phase boundaries summarized by the cartoon in panel **C**. Among the possible phases we highlight: an asynchronous state (♦), as well as oscillatory states with fast (•) or nested fast/slow (▪) frequencies. Panel **(B)** shows corresponding example activity traces. *See Figs. S1, S2 and S3 for additional information*.

For weak inhibition the dynamics settled in a regime characterized by constant levels of activity in all layers (representative traces are shown in Figure 2B, ♦ marker), as it would be observed in the case of asynchronous neuronal population firing with homogeneous rate. A rate instability line was crossed for stronger values of *K*_E_ beyond which firing rates diverged (white region at the top right of the dynamic regime profiles). The constant rate regime lost its stability as well for larger inhibition strengths. When the absolute value of *K*_I_ was gradually increased periodic oscillations emerged (traces in Figure 2B, • marker), in which all the layers oscillated with a fast frequency (in the gamma range) and similar relative amplitudes. When further increasing *K*_I_, more complex oscillatory patterns were observed (example traces in Fig. 2B ▪ marker, and Fig. S3). Among them were apparently chaotic and periodic oscillations distinguished by, respectively, low and high values of the autocorrelation’s highest peak (fourth row of Fig. 2A), and which appeared to be locked across layers at various frequency ratios as indicated by the delay of the autocorrelation’s first peak (third row of Fig. 2A).

Note that the parameters *K*_E_ and *K*_I_ are fixed for each given performed simulation. This is a quite artificial situation. In vivo, we can expect that under the influence of noisy drive, neuronal adaptation [44], short term plasticity [45], neuromodulation [46] and other causes the effective strength scales of excitation and inhibition slightly change over time and that, therefore, *K*_E_ and *K*_I_ fluctuate in the surroundings of an average working-point. In that sense, the complete dynamic regime profile is indicative of the diverse dynamical repertoire that the activity of a local circuit may sample at different times, especially if the working point is close to different phase boundaries, which can then be crossed by only slight changes of parameters (see Discussion).

For better visualization, we summarized all the qualitatively different dynamical modes in a cartoon regime profile, sketched in Figure 2C. Even more phase subdivisions could be generated by inspecting the relative phase of oscillation of the different layers. Before analyzing interlayer phase differences, however, we will first study the dominant frequencies of oscillations, which happen to be strongly dependent on the considered layers and dynamical regimes.

### Fast and slow oscillations dominate in superficial and deep layers

The oscillatory trace obtained for the working point marked by ▪ (referred to in the following as the “fast/slow working point”) in Fig. 2B clearly shows that spectrally rich oscillatory patterns can be generated by our model, simultaneously involving multiple faster and slower frequencies which are expressed more or less prominently depending on the considered layer. Notably, in the specific case of the fast/slow working point, L4 activity is dominated by a fast gamma-band oscillatory component around 71 Hz, which is also strong in L2/3, while lower layers L5 and L6 are clearly dominated by slower oscillatory frequencies in the alpha-/beta-range. This fact is intriguing, since as, previously mentioned, a similar segregation in the frequency of neuronal rhythms between superficial and deeper layers was observed also experimentally [24–27].

The pattern of layer-specificity of frequency shown by the fast/slow working point (▪) is not obtained by a fine-tuning of parameters, but represents a feature shared by an entire region in the *K*_E_-*K*_I_-plane, which we call the “fast/slow domain” and which is represented in dark blue color in Fig. 2C. For each point belonging to this region (see Materials and Methods for the calculation of its boundaries), we evaluated the relative amount of low power *P*_lo_/*P*_tot_ for each layer. As shown by Fig. 3A (computed for the same bottom-up input as in Fig. 2), within this region, deep layers’ activities—particularly L5’s—was generally dominated by slow oscillations, whereas L4 developed almost exclusively fast frequency oscillations. L2/3 manifested an intermediate behavior, with a balanced median value of *P*_lo_/*P*_tot_ across the whole region of approximately 50 %.

**Figure 3.**
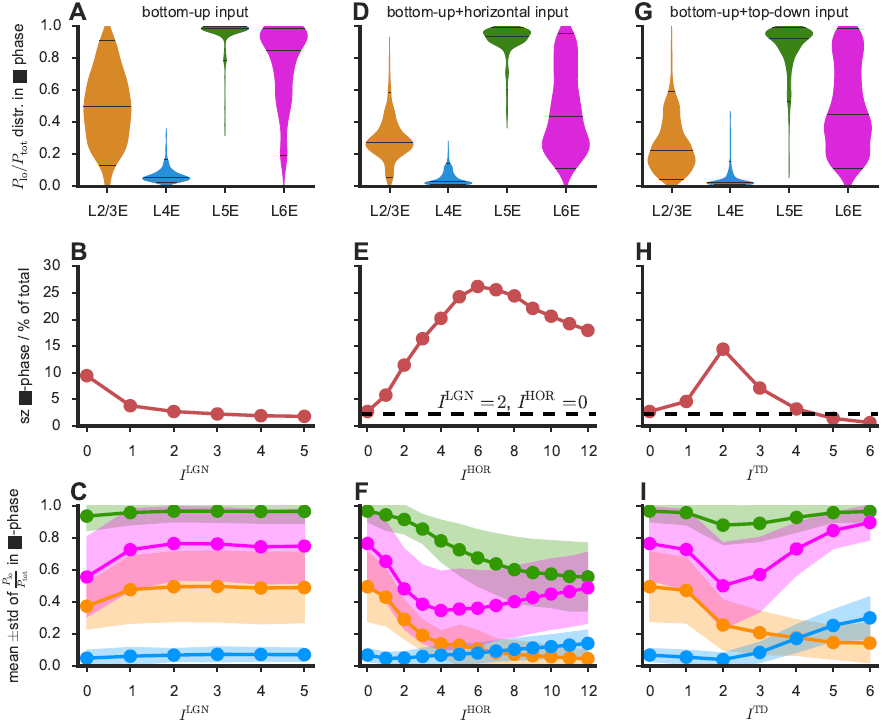
Dominance of fast and slow oscillations in superficial and deep layers, respectively. In the fast/slow regime (marked blue in Figs. 2C, S1C and S2C), L4 and to a lesser degree also L23 were dominated by faster oscillations (i. e. low fraction of low power *P*_lo_/*P*_tot_) when the local circuit was driven by bottom-up inputs only, as shown by the distributions in panel (**A**). Applying different combinations of contextual inputs (horizontal input to L23 in panel **(D)**, top-down input to L5 in panel **(G)**) modulated these distributions, further increasing e.g. the fast frequencies dominance in L23. This trend was also robust to changes in the stimulation strength (**C,F,I**), with a dependence on contextual inputs (**F,I**) more non-linear than on the bottom-up input (**C**). Likewise, the propensity to undergo such complex slow/fast oscillations (measured by the size of the fast/slow regime) was relatively unaffected by the strength of the bottom-up input (**B**), but peaked for specific horizontal and top-down stimulations (**E,H**).

The size and frequency distribution of the fast/slow domain depended on the specific context, i. e. the applied values of the bottom-up (*I*^LGN^), horizontal (*I*^HOR^) and top-down (*I*^TD^) inputs. In absence of horizontal or top-down drive, the size of this region was relatively constant for a wide range of contrasts, *I*^LGN^ (Fig. 3B), and for each of them the average distribution of high and low power over the layers (Fig. 3C) followed the same pattern as in Fig. 3A.

Dynamic regime profiles computed for different contexts (see Figs. S1 and S2 for applied horizontal and top-down input, respectively), showed that qualitatively similar regimes and equivalently rich dynamical repertoires continued to exist. Nevertheless, regime boundaries were distorted, producing variations of the extension and exact localization of the regimes summarized by Fig. 2C. The fast/slow region was very robust and continued to include the working point ▪ on the *K*_E_/*K*_I_ plane. The layer-dependent pattern of *P*_lo_/*P*_tot_, however, did change quantitatively. The distributions shown in Fig. 3A were altered into the ones displayed by Fig. 3D (by adding an horizontal input *I*^HOR^ = 2) or in Fig. 3G (by adding a top-down input *I*^TD^ = 2). In both these cases, the power balance of the more superficial L2/3 was shifted toward the faster frequencies of L4, while L5 continued to be dominated by slow frequencies. The dominance pattern of fast and slow frequencies was essentially maintained—despite some non-monotonic changes in L6—over the entire explored ranges of *I*^HOR^ and *I*^TD^, and the segregation of fast and slow frequencies in superficial and deep layers became even sharper for strong top-down inputs (see Figs. 3F,I). In addition, changes of horizontal and top-down drive also affected the size of the fast/slow phase. Unlike in the case of bottom-up drive *I*^LGN^, changes in size with *I*^HOR^ and *I*^TD^ were non monotonic, with the fast/slow-region reaching a maximum size for intermediate values of these contextual inputs.

Overall these analyses show that the fast/slow-region is robust—even actually enhanced—by the application of inputs related to contextual modulation.

### Dynamic phase leadership hierarchy between cortical layers

Besides variations in their spectral amplitudes at different frequency bands, the oscillations generated by our model also displayed a remarkable diversity of possible frequency-dependent interlayer phase-locking modes as a function of the strengths of excitation, inhibition and of context. To determine frequency-resolved oscillation phases we filtered each simulated time series in a narrow band around relevant frequency bands, such as the peak beta frequency in L23 (“L23E-*β* band”), or the peak gamma frequencies in L23 and L4 (“L23,4E-*γ* band”). After band-pass filtering, we identified oscillatory maxima in distinct cycles and, based on their time-stamps, we defined oscillation phase through a linear interpolation procedure. Details of the procedure for frequency-dependent phase extraction are provided in Materials and Methods (see Fig. S4A for a graphic summary).

We first considered the case of bottom-up drive only (*I*^LGN^ = 2, *I*^HOR^ = *I*^TD^ = 0). Fig. 4A shows dynamic profiles for the relative phase of L23 with respect to other layers, as a function of *K*_E_ and *K*_I_. We then focused on selected working points of interest. Choosing a working point very close to the transition between the constant rate- and the periodic oscillations-domain (*K*_E_ = 0.05, *K*_I_ = 0.65, marker **x**), all layers oscillated with a dominant fast frequency (cf. Fig. 2A and associated spectra in Fig. S4B). These fast oscillations were all *in-phase* locked, as revealed by the relative phase histograms of Fig. 4 (derived from oscillation maxima time-stamps in Fig. S4C).

**Figure 4.**
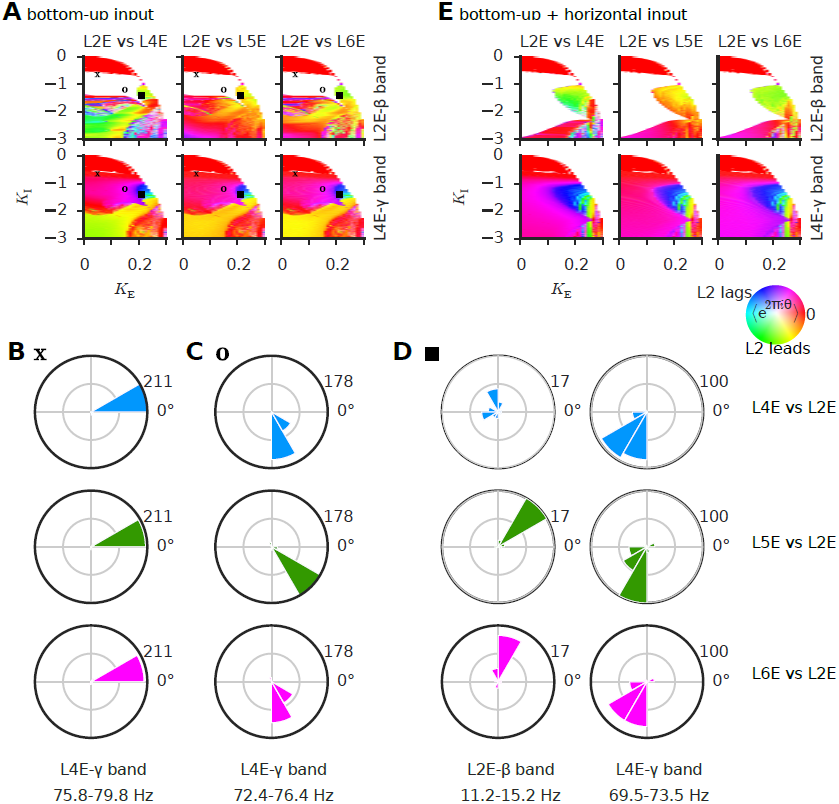
Frequency-dependent phase relations between cortical layers. **(A)** Regime profiles for average inter-layer phase differences show that multiple possible phase-locking patterns can be obtained in the model canonic local circuit. In particular, the region of periodic fast oscillations is subdivided into an in-phase locking (see **(B)** for phase difference distributions at the marker *x* working point) and an out-of-phase locking (see **(C)** for phase difference distributions at the marker *o*) sub-regions. In the fast/slow regime (including the fast/slow working point ▪) oscillations in layer 2 are leading those in the other layers for slow frequencies (**C**, left column), whereas the opposite was true for fast frequencies (**C**, right column). Similar phase difference configurations were robust toward contextual inputs, e.g. additional horizontal inputs to L23 (**E**). White background in panels **(A)** and (**E**) denote lack of appreciable oscillatory power in the corresponding frequency band. *See Fig. S4 for additional information.*

Moving further away from the homogeneous into the periodic oscillatory regime (*K*_E_ = 0.15, *K*_I_ = 1.2, marker **o**), oscillations of all layers remained fast but an *out-of-phase* locking pattern emerged in which L23 lagged behind L4 and the other layers (Fig. 4C and Fig. S4D). Thus, the fast-oscillating-regime (marked yellow in Fig. 2C) is actually subdivided into in- and out-of phase sub-regimes (Fig. 4A). This finding is in line with rigorous theoretical results about the phase-locking behavior of simpler models with symmetrically interacting oscillatory neuronal populations [47].

In the slow/fast-region (including the ▪-marked working point), phase relationships were different for the slow and the fast frequency bands of the observed multi-frequency rhythms. As evident from Figs. 4D and S4E, L23 was the phase-leading layer in the slow frequency band, but it lagged behind the other layers in the gamma band.

Addition of a horizontal input (*I*^LGN^ = 2, *I*^HOR^ = 2, *I*^td^ = 0), lead, as previously mentioned, to an expansion of the fast/slow region (Fig. 4E) with a largely similar phase-difference landscape.

### Dynamic phase leadership hierarchy between coupled local circuits in distinct cortical regions

Experimental evidence suggests that feedforward and feedback communication in the cortex might be subserved by fast and slow oscillations, respectively [28–32]. According to the communication-through-coherence hypothesis, in the case of out-of-phase locking, we expect that a stronger functional influence is exerted by the phase-leading onto the phase-lagging population, due to the fact that, in this direction, the target population receives the sent signals at a phase in which its excitability is elevated, favoring their transduction and integration [48, 49].

Given the functional relevance of dynamic phase-locking patterns in inter-areal communication, we then studied inter-layer phase relations when coupling together two canonic local circuits in the arrangement of Fig. 1F. Such an arrangement mimics in a simplified manner the layer specificity of cortico-cortical connections, giving rise to a structural hierarchy of cortical areas [21, 34, 35]. In this configuration we could distinguish a lower and an upper circuit module, meant to represent the canonic multi-layer structure of two cortical areas at different stages of the cortical hierarchy. Feedforward connections were assumed to proceed from population L23E of the lower circuit to populations L4E and L4I of the upper circuit. Feedback connections proceeded from L5E of the upper circuit to E and I populations in L23 and L5 of the lower circuit (for details see Materials and Methods). Concerning contextual inputs, the lower circuit received a bottom-up drive *I*^LGN^ to L4 and L6, as in the case of the isolated canonic local circuit, while the upper circuit received a top-down input current *I*^ATT^, identical for L23 and L5—i. e. identical to the joint application of *I*^HOR^ and *I*^TD^—, meant to represent a top-down signal activating attentional modulation by the upper onto the lower circuit [50, 51]. Note that a similar layer-specificity of attention-related signals was assumed in other models [18–20].

Phases and phase differences in Fig. 5 were determined as in Fig. 4, but instead of comparing time series of different layers in one column we now considered relative phases between matching layers in the lower and upper circuits (e.g. comparing L23 of the upper circuit with L23 of the lower circuit, etc.). Given its rich spectral properties, we focused on inter-circuit relations when the system was tuned to be at the fast/slow working point into the fast/slow phase (robustly preserved by the weak coupling of two circuits).

**Figure 5.**
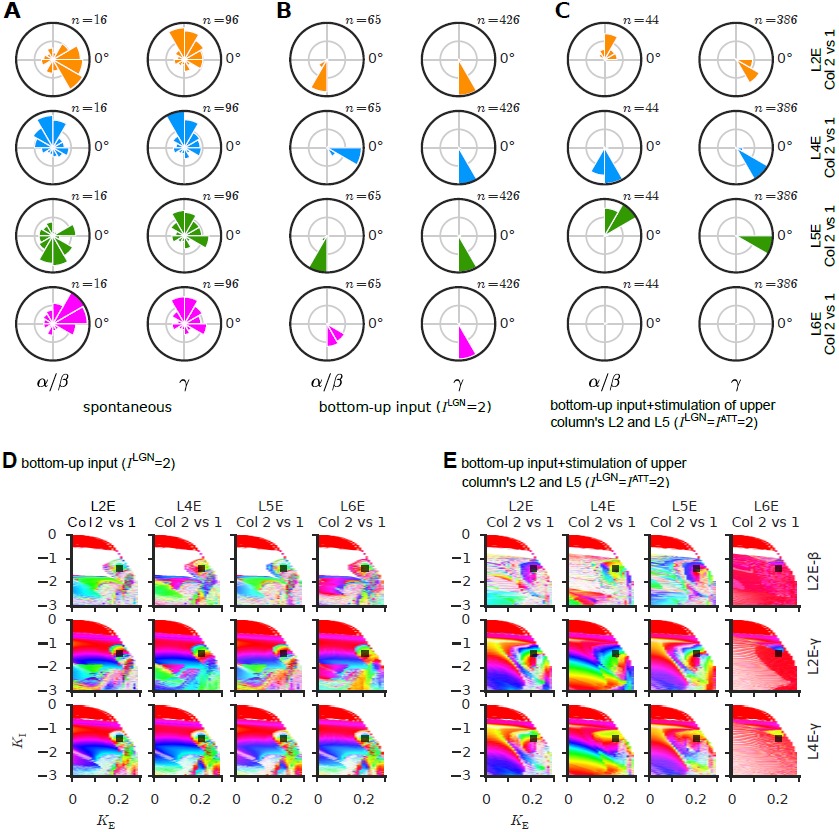
Frequency-dependent phase leadership hierarchy between connected cortical circuits. (**A-C**) Distributions of phase differences between homologous layers of two inter-connected canonic cortical circuits at different stages of the cortical hierarchy (cf. Fig. 1E) at the ▪-marked working point in fast/slow phase. (**A**) During spontaneous activity (background noise only), no specific phase locking between the different layers of the two circuits was found. (**B**) In presence of bottom-up input to the lower circuit, oscillations in the lower circuit lead in phase those in the upper circuit, consistent with a predominantly feedforward processing mode. (**C**) In presence of an attention-mimicking input (*I*^ATT^ inputs to upper circuit), the phase leadership hierarchy was frequency-dependent, with a feedforward processing mode in the gamma-band as before, but a feedback processing mode for layers 23 and 5 in the slow frequency band. Regime profiles for bottom-up input without (**D**) or in presence of (**E**) attentional modulation show robustly similar phase relations in the whole fast/slow phase. *See Fig. S5 for additional information.*

Without specific input, i.e. when the two-areal system was driven uniquely by background input *I*^BG^ = 1 to all layers, the weak long-range coupling strength adopted in our simulations was not enough to enforce stable phase relations between the circuits, and correspondingly, the distributions of relative phases between matching layers were broad and phase-locking often not significant (Figs. 5A and S5A). However, when the lower circuit was driven in addition by a bottom-up drive (*I*^LGN^ = 2), phase relations between the lower and the upper circuit became more tight, giving rise to out-of-phase configurations in which lower circuit layers—and in particular L2/3 of the lower circuit, the source of feedforward cortico-cortical projections—lead in phase over upper circuit oscillations (Figs. 5B and S5B), potentially favouring bottom-up inter-areal functional influences in both slow and fast frequency bands.

The situation changed in presence of emulated attentional modulation, when further contextual inputs *I*^ATT^ where applied to the upper circuit (Figs. 5C and S5C). In this case, while a bottom-up compliant phase-locking mode was maintained by all layers in the gamma-band, in the slow frequency band the upper circuit’s L2/3 and L5—the source of feedback cortico-cortical projections—were now leading over the matching layers in the lower circuit, in a way compliant with potentiated top-down functional influence of the upper onto the lower circuit at slow frequencies.

Beyond this specific example, evaluating dynamic profiles of the average relative phases (Figs. 5D,E) revealed that the entire phase including the considered fast/slow working point exhibited a similar dynamic pattern of phase locking between the columns (Figs. 5D,E).

### Layer-specificity of frequencies may stem from interlayer interactions

After characterizing the fast/slow phase and its robustness under variations of the input configuration, we investigated possible dynamical mechanisms that may explain its origin. In particular, we probed the causal role played by interlayer interactions. In order to determine their importance, we systematically tuned the strength of all excitatory and inhibitory interlayer couplings, by multiplying them with a factor Γ, varying between 0 (no interlayer connections) and 1 (default interlayer connection strength as above). We kept, on the other hand, the connectivities within each layer unaltered, in order to maintain intrinsic properties like their resonance frequencies.

We focused on the fast/slow working point (▪), in exclusive presence of bottom-up input (no contextual modulation). When all the layers were fully disconnected, (Γ = 0), L2/3 and L4 oscillated periodically at fast frequencies around 60 Hz and 70 Hz, respectively, as indicated by the harmonic peaks of the power spectra in Fig. 6A (top panel). On the other hand, L5 and L6 did not oscillate, as their intra-layer recurrent inhibitory interactions were not strong enough to destabilize the homogeneous fixed-point of activity via a Hopf bifurcation toward an oscillatory state [47]. Note that, with vanishing Γ and, therefore, in absence of the stabilizing effects of inter-layer inhibition, L6 activity tended to diverge, due to its complete lack of local recurrent inhibition, according to our chosen connectome [10], but most likely not in reality [52].

**Figure 6.**
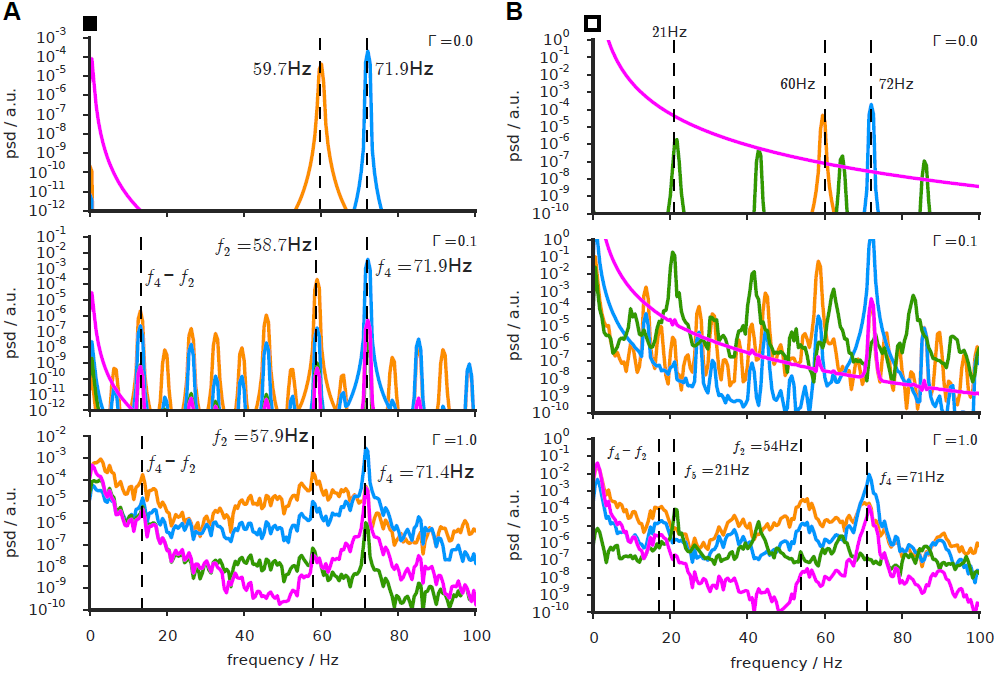
Layer-dependent fast/slow oscillations may be engendered by interactions between layers. **(A)** Power spectra of the oscillatory activity of different layers in the fast/slow working point, as a function of the strength of inter-layer coupling (modulated by a multiplier Γ). When layers are completely disconnected from each other (top panel, Γ = 0) L23 and L4 underwent intrinsically-generated periodic oscillations at slightly different fast frequencies *f*_4_ and *f*_2_. Deep layers did not intrinsically oscillate, but developed oscillatory activity already for a weak inter-layer connectivity (Γ = 0.1, middle panel). Appearance of new spectral peaks at frequencies such as the difference frequency *f*_4_ *f*_2_ indicates quasi-periodic oscillations. This peak and the intrinsic L23 and L4 peaks were still visible at full inter-layer coupling (Γ 1, bottom panel), even though spectra now developed an overall broad-band character, due to chaotic inter-layer entrainment. **(B)** For comparison, we show also power spectra for different inter-layer coupling strenghts in an analogous working point of a variant circuit model, in which properties of L5 have been modified to make it an intrinsic oscillator at a slow frequency *f*_5_. Gradual increase of inter-layer coupling revealed transitions in the structure of power spectra highly reminiscent of panel **(A)**, indicating that quasi-periodicity and chaotic entrainment subsist even in presence of intrinsically generated slow rhythms in L5. *See Figs. S6 and S7 for additional information.*

Once interlayer interactions were turned on (e.g. at Γ = 0.1, see Fig 6A, middle panel), power spectra became more complex, developing a large number of different peaks in all layers, among them peaks very near the original L2 and L4 fast frequencies, *f*_2_ and *f*_4_, observed at Γ = 0. We also observed peaks at a slower frequency (in the alpha/beta range) given by the difference of these two frequencies *f*_diff_ = *f*_4_ – *f*_2_ as well as harmonics of *f*_diff_, indicative of a quasi-periodic entrainment scenario [53]. Given that oscillations at slower frequencies such as *f*_diff_ could not be observed at Γ 0, we conclude that, in our simulations, such slower oscillatory components were caused by interlayer interactions, unlike the local intrinsic origin of fast oscillatory components within L2/3 and L4.

By further increasing the strength of these interaction up to Γ = 1 (Fig. 6A, bottom panel), the distributions of power spread to give rise to broad-band profiles, super-imposed with vestiges of the line peaks present at weaker Γ, which were more or less pronounced depending on the considered layer. In particular, the difference frequency *f*_diff_ was still recognizable in the spectrum which suggests the onset of chaos via quasi-periodicity [54]—as also observed in other, less detailed models of interacting oscillating neural populations [47, 48, 55].

This is further supported by bifurcation diagrams (Fig. S6B), in which the set of observed values of local maxima are plotted against the control parameter Γ, giving rise to tree plots whose increasingly complex branching structure describes the route to chaos (similar analyses have been performed in [47, 48]). Such bifurcation diagrams show characteristic windows of regularity, i.e. narrow ranges of Γ in which oscillations are regular and periodic (i. e. exhibiting only few discrete values for their local maxima). The spread of bifurcation trees for different layers, also reveal that chaos manifests itself more strongly in L23, than in L4, despite the mutuality of their interaction. This dynamical asymmetry may be interpreted as if L4 was effectively entraining L23 into chaos, in analogy with a purely feedforward entrainment case in which the oscillations of the driver population would remain perfectly unperturbed (cf. [48]).

While L4 power density was strongly concentrated in high frequency range for all interlayer coupling scales 0 ≤ Γ ≤ 1 (Fig. S6B, blue line), the relative fraction of low power grew markedly with Γ in layer 2/3 (Fig. S6A, orange line). Layers L5 and L6, which lacked an intrinsic fast frequency resonance, as previously discussed, became increasingly more integrated in the circuit as Γ grew from 0 to 1 such that at Γ = 1 their waveforms resembled a low-pass filtered version of L2/3’s (see Fig. 2B, and Fig. 6A, bottom panel), and, once they were sufficiently connected to the remaining circuit (Γ ⪆ 0.2), like L2/3 they also acquired increasingly more slow power as Γ grew towards 1 (Fig. S6A, green and pink lines).

Finally, the mutual entrainment dynamics between layers could be modulated by changes in contextual inputs. For instance, as shown in Figs. S6C–D, adding an increasing horizontal input stimulation to L2/3 (Fig. S6C) on top of the bottom-up drive to L4 leads to a situation in which the effective dynamic roles played by L4 and L23 are reversed. For strong horizontal stimulation, L23 is “strenghtened” and becomes the effective driver of L4. Correspondingly, L23 oscillations are more periodic and L4 oscillations more irregular, hence the altered relative fractions of low power, reported in Fig. 3F (in average over the entire fast/slow phase) and in Fig. S6D (in detail for the fast/slow working point ▪).

### Intrinsic and network-generated slow oscillations may co-exist in the same local circuit

Oscillatory components slower than high gamma frequencies emerged in our model thus as an effect of inter-layer interactions and disappeared when layers were disconnected (Fig. 6A). However, experimental studies in vitro showed that slow oscillations in deep layer can be pharmacologically induced even when infragranular layers are anatomically disconnected from supragranular ones [24] and that L5 is sufficient to promote synchronized activity at 1-12 Hz [56, 57]. In order to account for these findings that our original model cannot reproduce, we artificially modified its parameters ad hoc (see Materials and Methods) to make L5 intrinsically oscillating at a slow frequency. Doing so, we found that both intrinsic and network-generated slow oscillations may be simultaneously present and, therefore, hard to disentangle.

The intrinsic slow resonator model also gave rise to a rich dynamical repertoire, including a multiplicity of regimes for different choices of *K*_E_ or *K*_I_. Fig. S7 shows dynamical regime profiles, time-series from representative working points and a cartoon regime diagram for the modified model, analogously to Fig. 2 for the original model. Despite the appearance of additional regimes—such as a strictly periodic oscillatory regime encompassing both fast and slow frequencies in different layers (denoted by the M marker in Fig. S7B and C—, dynamical regimes reminiscent of those in the original model could still be recognized. In particular, there existed, at the same localization in parameter space as in Fig. 2, a regime of fast/slow oscillations. In analogy to the original model we analyzed the generation of these oscillations for a slightly different working point (denoted by □) through decoupling of the layers. Fig. 6B shows that, for this working point, oscillations continued to switch from periodic to chaotic when increasing the strength of the inter-layer coupling Γ. Due to the presence of an intrinsic slow resonance, when the layers were disconnected (Γ = 0) the power spectral density for L5 had a peak at around 21 Hz, incommensurate for our parameter choices with the intrinsic power peaks of L23 and L4. Nevertheless, increasing Γ, spectra became increasingly broader-band for all layers, as in absence of the L5’s intrinsic resonance (compare Fig. 6B with Fig. 6A). At Γ = 1, a vestige of the original intrinsic L5 peak at *f*_5_ 21 Hz remained only barely visible, differing merely by a few Hz from the *f*_4_ *f*_2_ peak whose origin lies in the quasi-periodic interaction with the other layers. Therefore, in presence of noise as in vivo, these peaks with very different mechanistic underpinnings—i.e., network interactions for the *f*_4_ – *f*_2_ peak, local intrinsic resonance for the *f*_5_ peak—may be difficult to distinguish.

Thus, dynamical mechanisms based on intrinsic layer-specific resonances and on inter-layer network-level interactions can coexist in the same model. Furthermore, these mechanisms necessarily interact such that attributing the origin of fast/slow oscillations to one or the other independent causes may appear questionable, as the behavior of a complex system cannot be reduced to its components’ behaviours (see *Discussion*).

### Layer-specificity of oscillation frequencies is robust again small changes of the connectome

The adoption of the connectome experimentally quantified by Binzegger et al. [10] naturally gave rise to a rich dynamical repertoire including a regime leading to the experimentally observed layer-specificity of fast and slow oscillations. We therefore investigated how tightly the occurrence of such a regime depended on choosing precisely this specific connectome. Indeed, despite the unprecedented quality of the connectivity exploration performed in [10], experimental uncertainty cannot be eliminated. Furthermore, other studies [9, 11, 58] reported different connectivity diagrams, not fully conforming to Binzegger et al’s. It is therefore important to analyze the robustness of the identified dynome against changes of the adopted connectome. We did so for the original model (i.e., without intrinsic slow resonator in L5) in presence of bottom-up input only, presented above in Fig. 2.

As a first step, we explored how the dynamical behavior of the model local circuit is affected by small changes of the strengths of its connections. For computational reasons, it was impossible to exhaustively explore the 64-dimensional space of disturbed matrices around the reference Binzegger matrix. We therefore probed only modifications along four selected directions of interest. To select these directions, we compared the interlayer connectivity matrices given by Binzegger et al. [10] and Haeusler and Maass [11] and identified four connection groups that were differing the most between these two connectomes, potentially hinting at a more pronounced experimental uncertainty in their determination. These connection groups were: inhibitory links with L23 (L2I ⟶ L2E and L2I ⟶ L2I, we termed this modification `mod22`); L4 inputs to excitatory population in L23 (L4E ⟶ L2E and L4I L2E, termed `mod42`); inhibition from L2 to excitatory population in L5 (L2I ⟶ L5E, termed `mod25`); finally, mutual recurrent inhibition in L5 (L5I L5I, termed `mod55`). We also modified simultaneously several of these connection groups, terming: `modX` the combination of `mod22, mod42` and `mod25`; and, (all) all four of these modifications together.

Given its functional relevance, we focused on how robustly preserved the segregation of fast and slow oscillation frequencies in superficial and deeper layers was. Therefore, we measured the relative area occupied by regions in the simulated dynamic regime profiles where the relative amounts of low power, *P*_lo_/*P*_tot_ were, respectively, greater than 50 % in deep (L5E, L6E) and smaller than 50 % in superficial (L2/3E, L4E) layers. The gradual changes of this relative area are plotted in Fig. 7 against the modification factor *α* along the different probed directions (see Materials and Methods for details). For the modifications `mod42, mod55` and `mod25`, as well as their combination modX, the relative extension of the regions showing segregation of fast and slow frequencies in superficial and deep layers, respectively, remained relatively constant at around 2.5 %, matching approximately the size of the fast/slow phase including the fast/slow working point ▪ in the undisturbed connectome (compare Fig. 3B).

**Figure 7.**
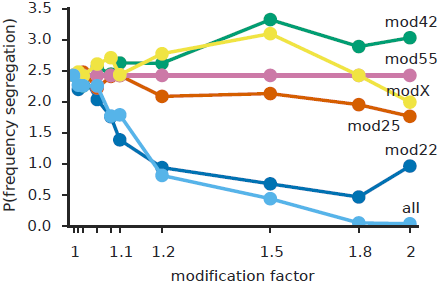
Layer-specific fast/slow oscillations are robust against small changes of the connectome. We modified the original connectome by scaling selected subsets of its connections by a continuously increasing modification factor. For the six patterns of modification we adopted (see Materials and Methods for details), the size of the fast/slow-region in the dynamic regime profiles remained nearly constant or reduced only gradually when increasing the modification factor, indicating that emergence of a fast/slow regime is a robust feature of the adopted connectome.

The system’s dynamics was more sensitive to the modification mod22, for which a morphing factor of *α* = 1.2 was sufficient to shrink the size of the occurrence of layer-segregated frequency-dominances by about 50 %. When applying all modifications at the same time (`all`) the segregation of fast and slow frequencies in superficial and deep layers was not anymore observed starting from *α* = 1.8. Nonetheless, also for the modifications `mod22` and `all`, sufficiently small morphing factors *α* preserved the frequency segregation. This indicates that this dynamical behavior is robust. It does occur when adopting precisely the Binzegger connectome, but also when assuming connectomes which lie in its *local neighbourhood* in the space of possible connectomes, i.e. which are obtained from it through a small continuous deformation.

### Layer-specificity of oscillation frequencies is compatible with multiple but non-random connectomes

As a second step, we explored the robustness of the layer-specificity of dominating frequencies against more substantial changes of the connectome. To do so we generated artificial connectomes, by randomizing all inter-layer connection strengths, but keeping all intra-layer connections unchanged in order to preserve each layer’s intrinsic properties. We analyzed the regime profiles of 100000 different randomized connectomes searching the *K*_E_ − *K*_I_ plane for a working point where all layers were oscillating, at least one of them predominantely fast, and at least one of them predominantely slow (see Materials and Methods for details). Remarkably, despite their not too stringent nature, these constraints were simultaneously satisfied by merely 125 of the 100000 tested randomized connectomes (Fig. 8A). Moreover, in almost all of these 125 “good” connectomes, the layer dominated by fast frequencies was superficial (L4 or L2/3) and the layer dominated by slow frequencies was deeper (L5 or L6), just as in the original Binzegger connectome and in agreement with experimental observations, probably because of the strong inhibitory self-loop within L4 and L2/3 making them intrinsically resonating at fast frequencies [47]. We can conclude, thus, that a slow/fast regime generally *did not* emerge starting from a random interlayer connectivity. On the contrary, its manifestation proved to be non trivial, requiring the selection of highly specific connectomes compatible with it.

**Figure 8.**
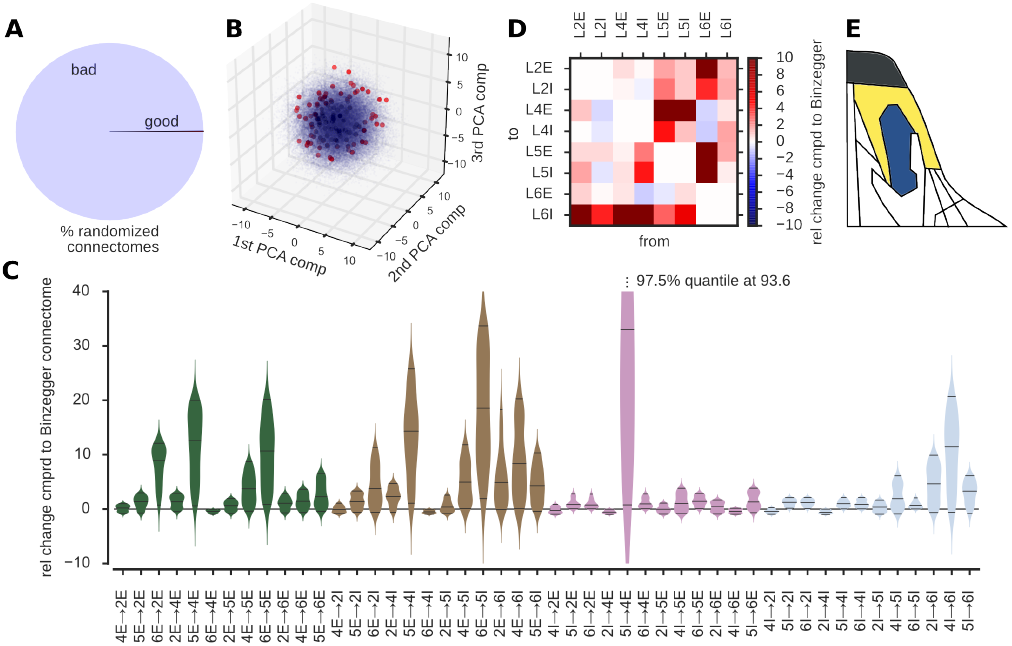
Non-random connectomes are required for the emergence of layer-specific fast/slow oscillations. We modified the original connectome more extensively by fully randomizing all its inter-layer connections. **(A)** Considering these randomized connectomes, we could identify a working point with layer-dependent fast/slow oscillations (akin to the ▪-marked working point considered above) in barely 0.1 % of the connectomes (termed “good”) that we tested. **(B)** Linear dimensionality reduction (PCA) did not reveal trivial features that could distinguish “good” (blue) from “bad” (red) connectomes. **(C)** There were substantial differences between alternative “good” connectomes, as shown by distributions of different connection weights over the set of retrieved good connectomes. **(D)** The connectivity matrix on an example randomized “good” connectome reveals clear differences with the experimentally determined reference one. Despite these differences, this connectome leads to dynamical regimes very similar to the ones of the original model, as indicated by a comparison between its cartoon phase diagram in panel **(E)** and the one of Fig. 2C. Matching colors are used for the asynchronous (black), the fast periodic (yellow) and the fast/slow (blue) oscillatory regimes. *See Fig. S8 for additional information and examples.*

Although very rare and thus, in this sense, non-random, there existed, on the other hand, more than just one “good” connectome. We therefore checked whether these “good” connectomes were structurally similar, i.e. whether they shared some specific structural feature, responsible for their overlapping dynamics. Given the small number of “good” connectomes, we expected the values of the different couplings to be highly interdependent. However, the precise patterns of interdependency proved hard to identify adopting conventional dimensionality reduction approaches. Principal component analysis (PCA) yielded no obvious separation of “good” from “bad” connectomes. Fig. 8B shows a scatter plot of the first three PCA components of the 100000 randomly generated connectomes. The highlighted “good” connectomes were distributed as raisins throughout the entire cloud of “bad” connectomes. Furthermore, a more detailed inspection of the distribution of PCA components, illustrated by Fig. S8A, failed to reveal any higher dimensional way of linearly separating “good” from “bad” connectomes. Beyond linear covariance decomposition, we attempted also other classification approaches—standard but more general and non-linear—, such as training of multi-layer perceptrons [59] or random forest regression [60]. Such approaches could correctly classify roughly 70% of the “good” connectomes used for training, but their generalization performance (tested via cross-validation loss) dropped to random levels. The identification of a much larger number of “good” connectomes would be required in order to improve the training and avoid over-fitting, but it may be computationally too demanding, given that in the order of 10^5^ connectomes needed to be simulated in order to obtain the dynamic regime profiles of in the order of 10^2^ “good” connectomes. In conclusion, thus, we were unable to identify any trivial common feature of those connectomes which possessed a slow/fast regime.

As a matter of fact, the strengths of many connections varied among the “good” connectomes in a (seemingly) unconstrained manner. Analyzing link by link the distributions of the various interlayer connection couplings over the “good” connectomes (Fig. 8C) we found that some excitatory connections, as well as the inhibitory connections to layer 6I and in particular the connection 5I ⟶ 4E could assume values over very wide ranges. Other links, on the other hand—mostly inhibitory connections with the noted exceptions—took on values within a much more restricted range, indicating a tendency not to deviate too far away from their measured value in Binzegger’s reference connectome. Fig. 8C also shows that connection strengths within the artificial “good” connectomes tended to assume higher values compared to the original Binzegger connectome. Moreover, the Frobenius norm (i. e. the squre root of the sum of squared entries) of the connectome matrices was higher for all randomized connectomes than for the original Binzegger connectome. While our randomization procedure might induce a bias towards higher matrix entries (as the entries are drawn from a uniform distribution, while the Binzegger connectome possesses relatively few strong, but many weak entries, see Materials and Methods for details) this could nevertheless hint to homeostatic plasticity mechanisms maintaining the cost in synaptic resources at a low level [61].

Fig. 8D (as well as Figs. S8C,E) illustrates exemplarily how pronounced differences between randomized “good” and the original Binzegger connectomes could be. Despite the difference in wiring, the associated regime diagrams still showed familiar features (see Fig. S8B for the regime profiles on which these phase diagram cartoons are based). In particular, as in Figs. 2A,C, we could clearly identify a rate instability line (right upper corner), an homogeneous regime (upper left corner) destabilizing into a periodic oscillation regime with all layers oscillating at fast frequencies (regime marked in yellow in Fig. 8E, and in Figs. S8D,F), and, although at a different location in the parameter space, the characteristic fast/slow oscillation regime (marked in blue in Fig. 8E, as well as in Figs. S8D,F), that was, by definition, required to be present for this “good” connectome. The dynome of the randomized connectome included also other regimes, not necessarily supported by the original Binzegger connectome, but we did not attempt matching them systematically, as they were not our main interest.

Finally, we coupled together to mimic hierarchical cortico-cortical interaction as in Figs. 1 and 5, two canonical circuits with the alternative “good” connectome of Fig. 8D. Interestingly, also in this case we could identify regimes with a frequency-dependent phase-leadership hierarchy, in which the upper circuit was phase-leading in slow frequency bands and the lower circuit in fast frequency bands (Fig. S8G). However, we did not study systematically how robust such phases are for the connectome of Fig. 8D, nor did we check whether similar regimes occur for all “good” connectomes or only for certain, thus further shrinking their number, when additional constraints are added (see Discussion).

## Discussion

Connectomics generally assumes that structural connections in the brain are crucial to constrain its function [62]. In line with this general tenet, our analysis indicated that simulations based on a realistic local circuit connectome could account for system states with high potential functional relevance. More specifically, we showed that a local cortical circuit embedding an experimentally determined multi-layer connectivity architecture robustly lead to a dynamical regime in which oscillations in the gamma- and alpha-/beta- frequency range occur predominantly in upper and lower layers, respectively, reminiscent of experimental findings [24, 26]. By coupling two of such circuits through laminar-specific cortico-cortical connections, in order to mimic interaction between different stages of the structural cortical hierarchy [21, 34], we found that the relative phases of neural oscillations in different circuit modules and layers spontaneously self-organized to favour feedforward communication-through-coherence in the gamma-band and—when a top-down signal (like the allocation of attention) was phenomenologically emulated—feedback communication-through-coherence in slower frequencies. Likewise, these results are compatible with empirical evidence [30, 32]. Last but not least, we established that random connectomes would lead to qualitatively different dynamical behaviors, confirming from a dynamical perspective the importance of gathering high-quality structural connectivity data [63] and of utilizing them in theoretical or computational work [64].

Beyond a linear one-to-one mapping between structure and function, we stress nevertheless that a given fixed structural topology can give rise to a multiplicity of dynamical regimes and thereby to very different functional connectivities [48, 65], which is not surprising from a dynamical systems perspective. When the working point of the local circuit is chosen to give rise to to be in vicinity of phase transition boundaries, noisy fluctuations or weak modulatory biases may thus easily trigger qualitative modifications of circuit behavior in absence of any structural connectivity change. For instance, the analyses of Fig. 2 indicate that, for our deterministic model, at least three phases lie within very narrow ranges of the effective strengths of excitation and inhibition: the homogeneous phase (♦ marker); the periodic phase (• marker), with multiple possible variants of inter-layer phase relations; and, the fast/slow phase (▪ marker) on which our study chiefly focused. Selecting a working point in these parameter ranges may hus lead to a rich in vivo dynamics in which asynchronous firing alternates, as an effect of fluctuations, with spectrally complex oscillatory epochs, that have layer-specific dominant frequencies and give rise to meta-stable transient phase-locking. Note that working points that are close to all the three aforementioned phases, would also lie at the edge of a rate instability line (at the upper right of Fig. 2C). This feature has been associated with non-linear amplification capabilities, useful for stimulus selection [66], or to the occurrence of neuronal avalanching behavior [67], even if the simplifications inherent to our model (see later) do not allow us to deal with these phenomena in our study.

A fixed structural connectome can not only give rise to switching between multiple functional states (“functional multiplicity” [68]), but our study also illustrates the complementary case: multiple well distinct structural connectomes engendering analogous functional states (“structural degeneracy” [68]). Through a non exhaustive search we determined that roughly a hundredth of all tested random inter-layer wiring diagrams are compatible with the existence of a network-generated fast/slow oscillatory regime. These connectomes could have largely equivalent regime profiles, but we have also shown that they do not share any obvious structural feature responsible for their common dynamical behaviour. The failure of linear dimensionality reduction approaches such as PCA indicates that any criterion for predicting that a connectome will be “good” would necessarily be a non-linear function of a high-dimensional vector of connectivity parameters. More general and non-linear classification approaches such as training of multi-layer perceptrons [59] or random forest regression [60] may be used to learn this criterion from brute-force extracted samples of “good” connectomes. Nevertheless, our preliminary attempts to do so have failed to achieve generalization performances above chance level. Larger training sets would then be needed to extract predictive information via machine-learning approaches, but obtaining them may be computationally too demanding, given that to identify *O* (10^2^) “good” connectomes we already had to perform a systematic parameter exploration for *O* (10^5^) randomized connectomes.

The fact that it is possible to artificially build different connectomes with similar dynamical properties does not necessarily imply that such alternative connectomes may actually be implemented somewhere in the brain. Many studies indeed advocate the existence of a unique canonic local circuit repeated with minimal variants throughout the entire cortex [5, 7, 69–72]. On the other hand, this view has also been criticized, because of large differences in local circuits found between areas and species [73–75]. Other studies proposed that multiple canonic circuit types may exist and be adopted by different brain areas [76, 77]. However, it is not clear whether these differences in structure correspond to actual differences in function [78]. Our computational approach suggests that both small local modifications of the “standard” canonic local circuit (Fig. 7), and suitable global rewirings transforming it into one of the alternative “good” connectomes may have a limited impact on the emergent dynamical repertoire. Unfortunately, we still miss a systematic charting of local circuit variants in the cortex. For the moment it is therefore impossible to verify to which extent some of our synthetically generated “good” connectomes match existing cortical circuits, although experimental information to attempt this task may become available within the next few years, thanks to ongoing large-scale efforts [79].

We defined a connectome as “good” only in virtue of its capacity to give rise to a fast/slow oscillatory regime. Obviously, however, we do not expect this capacity to be the unique purpose of the canonic local cortical circuit(s). It is possible that when additional target functions are prescribed, the actual number of “good” connectomes drops even more, as an effect of additionally imposed constraints. Structural degeneracy may even vanish, and a single, or just a few, canonic circuits be retained when the true or, at least, a more complete ensemble of criteria to be fulfilled is assumed. Another possibility is that many circuits may implement the same dynamical and functional repertoire, but that only a few of them have been actually selected during evolution or development because they are, in addition, optimal in some sense (wiring length, communication efficiency, compliance with developmental chronology, etc.) [80–83]. A potential hint to the presence of an optimization mechanism akin to developmental homeostatic plasticity [84] may be the small overall cumulative synaptic weight of the reference connectome derived from [10] compared to the other artificially generated connectomes. Finally, a third scenario is that structural degeneracy subsists even when the number of constraints is increased, because—beyond mere redundancy—it is useful from a system’s perspective, conferring additional robustness to some canonic set of essential functions against unexpected perturbations or ecological changes [85]. Such a scenario is known for instance to occur in invertebrate neural systems devoted to vital functions [36–38, 86]. Again, more systematic connectivity reconstructions are required in order to substantiate one or the other conjecture.

At the present stage, we can only acknowledge that the local cortical circuit wiring diagram of Fig. 1—as well as the alternative connectomes of Figs. 8D and many others—play a key role in the genesis of non-trivial dynamical regimes such as the fast/slow oscillatory phase, but we cannot fully explain why this is the case. Our anlyses have nevertheless revealed that, in our model, fast oscillations generated intrinsically within specific layers engender slower oscillatory components. Note that, as discussed in relation with Fig. 6B and Fig. S7, different mechanisms for the generation of slow oscillations may coexist. In particular inter-layer entrainment is not incompatible with the existence of intrinsic slow-frequency resonances in infragranular layers [24, 33, 56, 57]. On the contrary, the fact that the collective activity of all neuronal populations in these layers already tends to develop slow oscillations due to entrainment may favour the local recruitment of interneuronal populations intrinsically resonating at these slow frequencies. Our model predicts that the inter-layer entrainment dynamics can be biased by changes in contextual inputs or stimulation of specific layers (cf. Fig. S6) and this prediction may allow to probe the presence of inter-layer entrainment in future experiments, by checking whether “devil’s staircase”-like transitions in oscillatory properties, such as frequency and power ratios, are induced when a control parameter is smoothly varied (cf. discontinuous steps in the *P*_lo_/*P*_tot_ curves of Fig. S6A and C).

Interactions between layers may also be interrogated optogenetically through selective stimulation of specific neuronal populations in specific layers [87–89]. Nevertheless, different experiments often yield incompatible results and the direct interpretation of stimulation effects in terms of circuit mechanisms has proven difficult [90]. The extreme simplifications adopted in our model—such as a collective neural mass description of neuronal populations ignoring neuronal diversity and specific micro-connectivity and timing patterns [91–93]—may cast doubt on the reliability of quantitative predictions based on it. Nevertheless, from a qualitative perspective our study anticipates that the effects of selective stimulation would strongly depend on the actual working point of the system. Furthermore these effects would be complex and not limited to simple transient and local changes of the firing rate, but also affecting phase relations, power spectra of oscillations in different layers, stability of entire dynamical regimes, and other properties at the system level (cf. Figs. 3, 4, S1 and S2).

Several models before ours have explored how multi-layer connectivity shapes dynamics. A previous study from our group [55] already investigated the impact of inter-layer connections on induced cortical oscillatory activity, linking the emergence of broad-band oscillatory spectra to inter-layer entrainment. Noteworthy, this earlier model involved interacting populations of spiking neurons, rather than rate units and included a very simplified layer structure, reduced to just two layers, thus confirming the robustness of the effective entrainment dynamical phenomenon against changes of the model architecture.

Other independent studies incorporated a more realistic layer structure. For instance, Binzegger and colleagues implemented a computational rate model of a local cortical circuit [66] based on their experimental anatomical reconstructions. This study conjectured that the circuit working point must be close to a rate instability, as we also suggest, but did not explore the possibility of functional multiplicity nor the generation of oscillatory behavior.

Wagatsuma, Potjans and colleagues studied the dynamics of one or multiple coupled canonic local circuits, performing efficient and detailed large-scale spiking network simulations and phenomenologically modelling the action of attention [18–20]. Their model gave rise to a regime of activity, characterized by realistic, generally very low, spontaneous firing rates for all layers, which increase from superficial to deeper layers with the exception of L6E. In comparison, firing rates in our model also followed a similar pattern (cf. Fig. 2A, first row), except that firing rates in L6E were much higher. Nevertheless, such discrepancy could be ascribed to the (potentially erroneous) lack of feedback connections from L6I to L6E in our connectome inspired from [10], unlike in the connectivity scheme in [18–20], where anatomical information from additional sources was used [58]. Furthermore, Wagatsuma, Potjans and colleagues described effects of attention on single layer firing rate, i.e. a reduction in L4 and an increase in L23 and L5. Note that similar antagonistic variations of rate in different layers can be caused by contextual modulations even in our model, but in a way which depends on the chosen working point (compare Fig. 2A with Fig. S1A and S2A). However, as [66] before them, Wagatsuma, Potjans and colleagues also ignored the possible generation of synchronous oscillations, focusing instead on an asynchronous regime.

Besides, [55], only few modelling works explicitly accounting for cortical layer structure addressed oscillatory behavior, and if so, they either did not find substantial inter-layer differences in this behavior [94], or did not perform layer-specific spectral analyses [95]. The present study intends to amend this gap, reporting a rich repertoire of different possible oscillatory configurations, with different layer-dependent power spectra as well as local and long-range phase-locking profiles. Nevertheless all these configurations are stable states in our model, giving rise to sustained oscillations. They thus correspond to an elevated level of synchrony which is certainly unrealistic with respect to in vivo oscillations that are stochastically bursting and short-lived (see, e.g. [96]). Nevertheless, we expect that the stable oscillatory states of our deterministic rate model provide a good hint about the properties of the meta-stable oscillatory transients that a corresponding noise-driven spiking model would generate, when tuned to be at the edge of developing oscillatory synchrony [47, 55, 97, 98].

Models by Haeusler and colleagues [11, 99] finally focused on yet another perspective, finding that the canonic local circuit structure is beneficial for local computations, like e.g. pattern classification of current and past inputs. Our rate model is unfit to address questions linked to spike-based local information processing. Our perspective interest goes rather toward the modelling of self-organized information routing, given that layer structure shapes out-of-phase phase-locked oscillations in a way which is likely to profoundly impact on the efficiency of communication between neuronal populations [49]. Indeed, it has already been shown that changes in phase relations between collective population oscillations affect the routing of information codewords conveyed by spike patterns [48, 98, 100]. Our study suggests thus that the canonic local circuit structure—or structures, taking into account the possibility of structural degeneracy—is particularly fit to to implement communication-through-coherence simultaneously across multiple frequency bands, in contrast to local computations, for which random wiring architectures may also performing equally well as connectome-based circuits [101].

Ultimately and naturally, our multi-layer model could be adopted as a neural mass describing the regional activity of a brain region embedded within a larger-scale mean-field model of thalamo-cortical networks [102–104]—incorporating weighted connectivity information about feedforward and feedback cortico-cortical projections, as well other possibly relevant pathways, such as the laminar-specific connections with the pulvinar [105]—in order to reproduce the emergence of brain-wide multi-frequency coherence networks [31, 106, 107], to probe the determinants of their alteration in pathological conditions [108] and to understand how their dynamics might be controlled [109, 110].

## Materials and Methods

### Model

We simulate the rate response *r*_*kα*_ of a population *α ϵ* {E (xcitatory), I (nhibitory)} in layer *k ϵ* {1, 2&3, 4, 5, 6}. Each layer receives recurrent inputs with a delay *D* = 0.1 time units from within the column. The strength of these inputs, denoting the total charge induced into the postsynaptic population *lβ ϵ* (*k* {1, 2&3, 4, 5, 6}, *β ϵ* {E, I}) over all existing connections *lβ ⟶ kα* is written as 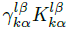 where 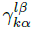 is the fraction of all synapses from population *β* to population *α* that are formed between layers *l* and *k*; values for 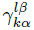 were taken from Fig. 12 in [10]. 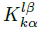 then gives the total number of synapses between populatoins *β* and *α* times the average charge induced in the postynaptic population due to a single spike in the presynaptic population. We decided to use 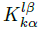 as parameters and explore the influence of these parameters on the behavior of the model. To make this feasible, we made the simplifying assumption that 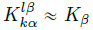, thus reducing the effective number of parameters to two, *K*_E_ and *K*_I_.

Apart from recurrent input the column is driven by four kinds of constant external currents. All layers receive the same background current *I*^bg^ modeling diffuse neuronal noise. Bottom-up stimuli influence the model column via currents *I*^LGN^, *I*^LGN^/3 and *I*^LGN^/6 sent to layers L4E&I, L6E and L6I, respectively, in accordance with [10]; the strength of *I*^LGN^ can be interpreted as the contrast of the stimulus. Contextual influences on the column are divided into horizontal, *I*^hor^, and top-down, *I*^td^, currents, targeting layer L2/3E&I and L5E&I, respectively, as suggested by experimental evidence [**?**]. A cartoon illustrating the complete circuit is shown in 1.

The total input into each layer, *I*^tot^, from both intrinsic and extrinsic sources is is assumed to activate the layer, if completely uncoupled from the others and after transients have settled, to a level given by the *f* − *I*-curve

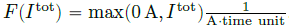. In summary, our model column was governed by the equations

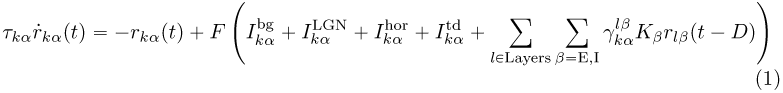

For simplicity, the relaxation time constants *τ*_*kα*_ were taken to be 1 time unit for all *k* and *α*.

When considering two coupled columns we extended that model by introducing feedforward connections from the lower column’s layer 2E to the upper column’s layer 4 (both the excitatory and inhibitory population), and feedback connections from the upper column’s layer 5E to the lower column’s layer 2 and 5 (both the excitatory and inhibitory populations). The complete equations read

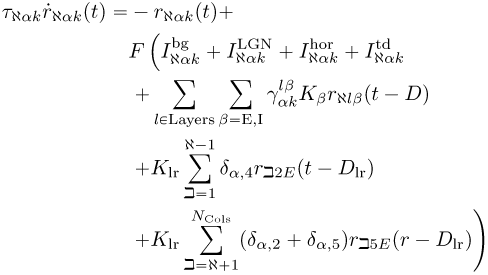

where *N*_cols_ = 2, 1 ≤*N,* ⊐ ≤ *N*_cols_ denote columns (which are ordered ascendingly according to their position in the cortical hierarchy), *δ*_*α,x*_ is the Kronecker-symbol, *K*_lr_ scales the efficacy of the long-range connections (we used *K*_lr_ = 1), *D*_lr_ is the delay of long-range connections (we chose *D*_lr_ = 10*D*) and all remaining parameters are as previously described.

The rate equations have not specifically been obtained through a mean-field reduction, but they do resemble a simplified Wilson-Cowan equation [111]. They were integrated with a custom written C code using a Runge-Kutta algorithm with time step 0.0001 time units. To overcome initial transients the first 200 time units (i. e. 2 000 000 time steps) were discarded and data anslysis was performed on the subsequent 2^20^ time steps. Deviating from that, Figs. 4B,D,F are calculated from time series with 2^21^ time steps. As initial condition for the delay differential equation we assumed that all time series were 0 for times *t* ≤ 0.

### Data analysis

All data analysis was performed in Python. Power spectra were calculated using Welch’s method with a window size of 2^19^ time steps, a window overlap of 2^18^ time steps and with the time series’ means subtracted. To make frequencies concrete, we set the transmission delay *D* to 10/3 msec, i.e. the time unit to 10*D* = 1/30 sec, i.e. the sampling frequency to 3 10^5^ Hz. Therewith, we defined low power *P*_lo_ as the summed power spectral density between 0 Hz and 30 Hz, and the total power *P*_tot_ as the sum of the power spectrum over all frequencies.

Cross-correlations were estimated by 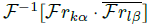 where *Ƒ* denotes the fast Fourier transform, *Ƒ*^-1^ the inverse fast Fourier transform, and 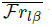 the complex conjugate of *Ƒr*_*lβ*_.

Cartoon regime profiles were obtained by semi-transparently overlying all layers’ profiles for the rate, fraction of low power, first delay and highest value of the autocorrelation and then tracing “obvious” boundaries. While some judgment calls had to be made this procedure is in no way critical as it only serves to highlight qualitative differences and similarities of the dynamics for different values of *K*_E,I_.

To assess the extent of the slow/fast-region we determined those points in the dynamic response profile that are “neighbors” of the point (*K*_E_, *K*_I_) = (0.24, *–*1.4), neighbors of these neighbors, and so on. For that matter, a point on the grid of sampled *K*_E,I_ values, reachable with one step either orthogonally or diagonally from a given source point is called a “neighbor” if it fulfills the condition that the fraction of low power was smaller than 0.999999 in all layers and greater or equal than 0.6 in at least one layer. We checked that the (*K*_E_, *K*_I_) points selected by this operational definition correspond to the visible outline of the slow/fast regime in the dynamic regime profiles.

### Oscillation phases

Fig. S4A illustrates how relative phases for oscillations of a predominant frequency are calculated: The power spectrum was estimated with Welch’s method as described above, then the global maxima of the power spectral density of layer 2 in the frequency band 8-30 Hz, of layer 2 in the frequency band 30-90 Hz, and of layer 4 in the frequency band 30-90 Hz were determined, and for each of these frequencies the time series of all populations, subsampled by a factor of 256 for performance reasons, were filtered with a finite-impulse-response filter, using a Kaiser-window of length 2^10^ with parameter *β* = 3.5 and a pass-band of ±2 Hz around the peak frequencies.

To determine the relative phase between the oscillations in population *a* relative to population *b*, local maxima were detected in the filtered time series of these populations. Assume for the moment that population *b* possesses at most as many local maxima as population *a*. For each local maximum 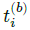 of population *b* we then located the times 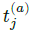 and 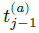 of the closest local maximum of population *a* occurring after and before 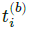, respectively. Let 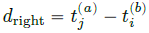 and 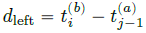. Therewith, we defined the relative phase as *d*_right_/(*d*_left_ + *d*_right_) and the phase-locking index *ϕ* as *ϕ ⟨*exp (2*πid*_right_/(*d*_left_ + *d*_right_)) ⟩_*i*_, where ⟨ ·⟩_*i*_ denotes averaging over all local maxima of *b*. In case population *b* had more local maxima than population *a* we used 1 – *ϕ*.

Note that while we usually used 2^20^ time steps (after the initial transients) for each time series, the right panels of Figs. 4B,D,F are calculated from time series with 2^21^ time steps.

### Alternative model with slow intrinsic oscillator in L5

To introduce a slow intrinsic oscillator into layer 5 we multiplied the matrix elements 5*I ⟶* 5*E* and 5*I ⟶* 5*I* in the original Binzegger connectome [10] by a factor of 400. When all interlayer connections were then set to 0 (i. e. Γ = 0) layer 5 could exhibit a peak in its power spectral density that was absent without this modification (see Figs. 6A,B top panels). Moreover, for full interlayer coupling (Γ = 1) and at least under bottom-up input (*I*^LGN^ = 2) L5 was then oscillating at predominantely slow (i. e. smaller than 30 Hz) frequencies as long as *K*_I_ ⪅ – 0.8 (see Fig. S7A).

### Robustness of results against small changes of connectome

We compared the connectome from [10, Fig. 12] to the one from [11, Fig. 1] by normalizing each separately with its respective maximum value and looking for the six biggest absolute values in the difference which were divided into four groups. The matrix elements of each group were then modified in order to determine their effects on the dynamics of the model column. In summary, we used the following manipulations: first, we changed the couplings *C*_L2I⟶L2E_ ⟶ *C*_L2I⟶L2E_/*α* and *C*_L2I⟶L2I_ ⟶ *C*_L2I⟶L2I_{*α* (we termed this modification mod22); second, *C*_L4E⟶L2E_ ⟶ *αC*_L4E⟶L2E_ and *C*_L4I⟶L2E_ ⟶ *αC*_L4I⟶L2E_ (mod42); third *C*_L2I⟶L5E_ ⟶ *αC*_L2I⟶L5E_ (mod25); fourth, *C*_L5I⟶L5I_ *αC*_L5I⟶L5I_ (mod55); fifth, the last three of these modifications together (modX); and sixth, all of the first four modifications at the same time (all); we run simulations for all of these six modifications for values *α* ϵ {1, 1.01, 1.025, 1.05, 1.125, 1.5, 2} on a grid *K*_E_ = 0 *…* 0.3 and *K*_I_ = –3 *…* 0 with stepsize Δ*K*_E,I_ = 0.01 and assessed how the fraction of working points p*K*_E_, *K*_I_q (excluding those with diverging rates *r*_*kα*_) where the relative amount of low power *P*_lo_/*P*_tot_ was above and below 50 % in superficial and deep layers, respectively, changed with *α*.

### Randomized connectomes

We randomized the Binzegger connectome to determine its relevance. To that end each connection was assigned a random value from a uniform distribution that was specific to the type of connection, *E* ⟶ *E*, *E* ⟶ *I*, *I* ⟶ *E* or *I* ⟶ *I*. The lower and upper bounds of these distributions were the minimum and maximum values of all matrix elements in the original Binzegger connection matrix belonging to the resepective connection type. Connections to and from layer 1, as well as those within a layer were completely disregarded in this procedure.

For each randomized connectome we simulated bottom-up input (*I*^LGN^ = 2, *I*^hor^ = *I*^td^ = 0) for a grid of working points, *K*_E_ = 1/30 *…* 1, *K*_I_ = –10 *…* 0 with a step size of Δ*K*_E,I_ = 1/30 which was traversed in order of increasing distance to the reference working point (*K*_E_, *K*_I_) = (0.21,*–* 1.4). To speed up the search, we first run a coarse-grained simulation for each working point, with a temporal step size of 0.001 and analzed 2^16^ time steps (after the initial transients). If total power *P*_tot_ in all layers was above 10^−5^ and there existed at least one layer with more than 25 %, and at least one other layer with less than 75 % of relative amount of low power *P*_lo_/*P*_tot_, we run another simulation for the same working point with the same, more fine-grained parameters that we used in the rest of the analyses, that is with a time step of 0.0001 and with 2^20^ time steps (after the initial transients). If we found that *P*_tot_ > 10^−5^ in all layers, and that there existed at least one layer with more, and at least one other layer with less than 75 % of *P*_lo_/*P*_tot_ we stopped traversing the grid of working points for the given connectome and termed it “good”.

### Figure details

In Figs. 2, S1, S2 and S3 used working points, defined by the tuple *K*_E_, *K*_I_, were ♦ = (0.05, *–*0.3), • = (0.1, *–*1) and ▪ = (0.21,*–*1.4), ▾ = (0.05,*–*1.7), = (0.05, *–*2), ◂ = (0.05, *–*2.5), ▸ = (0.13,*–*1.8), (♦ = (0.24,*–*2.3), • = (0.28, *–*2.9), • = (0.225,*–*2.9). In Fig. 2B panels show excerpts of 50000 (♦, •) and 200000 (▪) time steps.

In Fig. 3, bottom-up input (panels A) was simulated with *I*^LGN^ = 2, bottom-up+horizontal input (panels B) with *I*^LGN^ = *I*^HOR^ “ 2, and bottom-up+top-down input (panels C) with *I*^LGN^ = *I*^TD^ = 2.

In Fig 4, used working points (*K*_E_, *K*_I_) were x = (0.05, –0.65), o = (0.15, –1.2), ▪ = (0.21, –1.4).

In Figs. 6 and S7, used working points, defined by the tuple (*K*_E_, *K*_I_), were • = (0.1, –0.5), ◯ = (0.1, –1), □ = (0.22, –1.4)

## Acknowledgments

This research was supported by the Bernstein Center of Computational Neuroscience Göttingen (grants 01GQ0433, 01GQ1005B and 01GQ1005C). DB was supported by the Marie Curie career development fellowship FP7-IEF 330792 (“DynViB”).

## Supporting Figures

Figures S1 to S8.

**Figure 9 (following page). [Figure S1].**
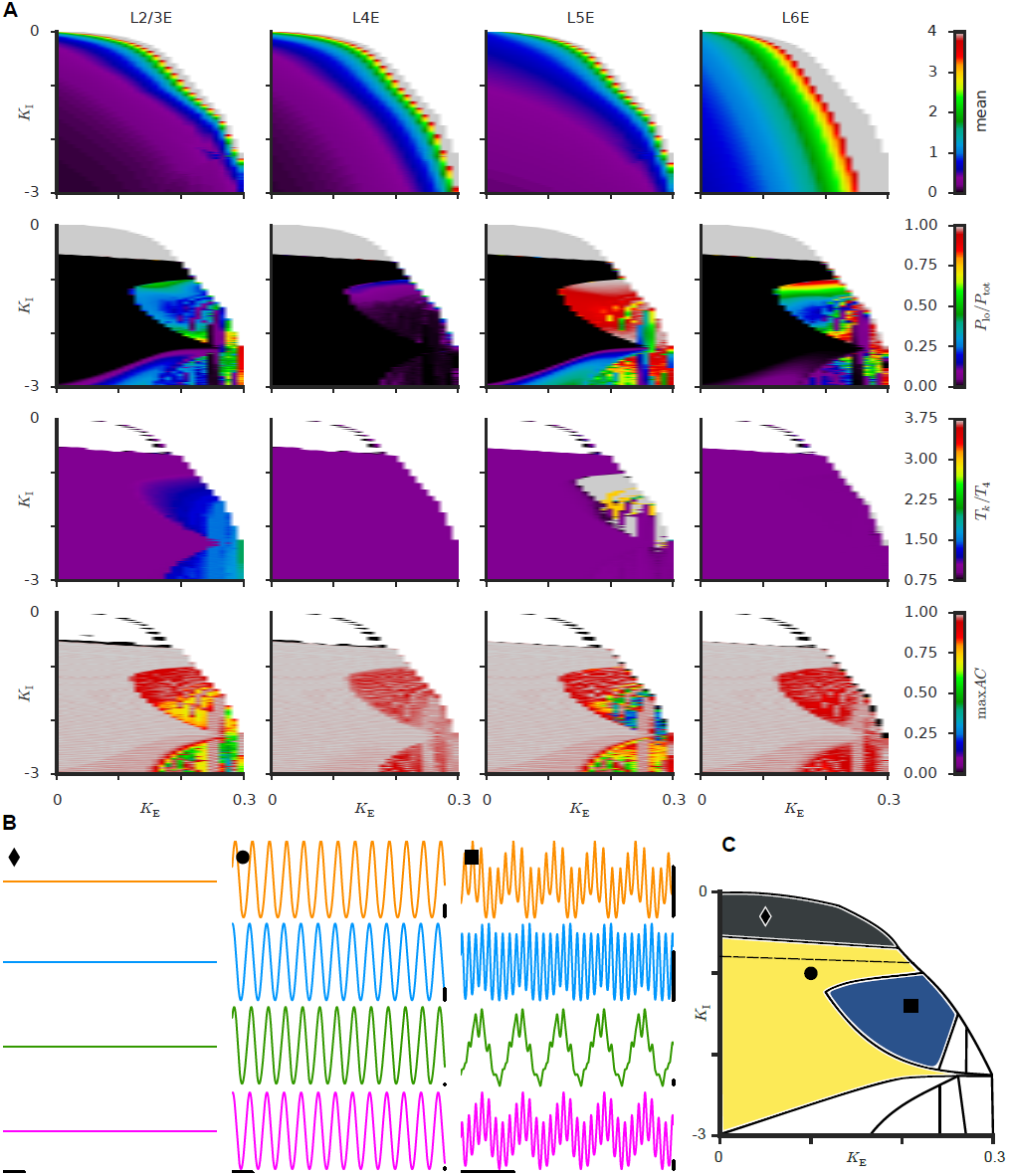
Dynamical regime profiles under bottom-up and horizontal (i.e. to L23) stimulation. All panels analogous to Fig. 2, but with additional input from horizontal connections to L23, in addition to bottom-up drive (*I*^LGN^ = *I*^HOR^ = 2, *I*^TD^ = 0). Asynchronous, all-fast and mixed fast and slow oscillatory phase are still visible, and are marked in panel C with the same color (dark gray, yellow and dark blue, respectively) as in Fig. 2C.

**Figure 10 (following page). [Figure S2].**
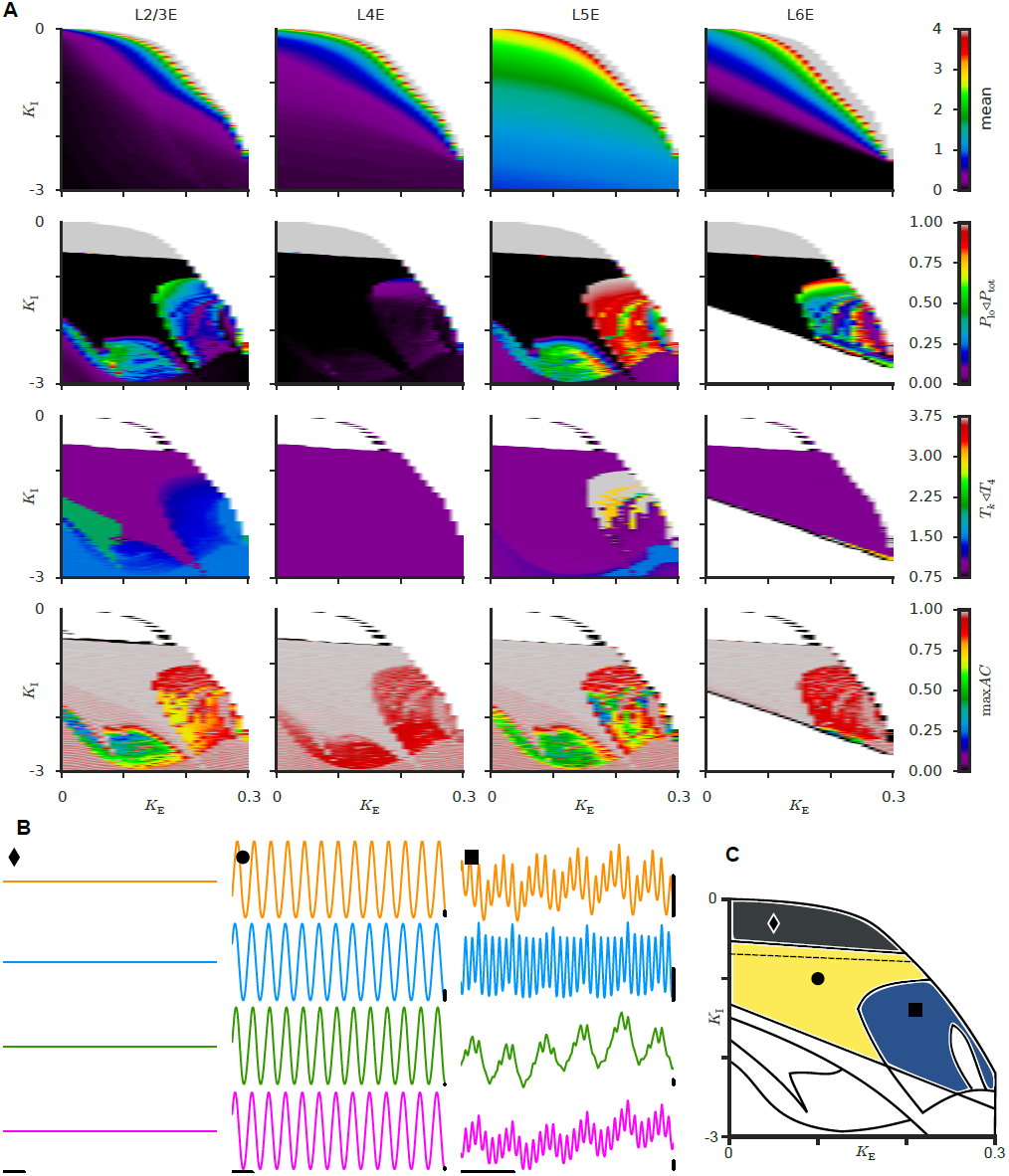
Dynamical regime profiles under bottom-up and top-down (i.e. to L5) stimulation. All panels analogous to Fig. 2, but with additional input from top-down connections to L5, in addition to bottom-up drive (*I*^LGN^ = *I*^TD^ = 2, *I*^HOR^ = 0). Asynchronous, all-fast and mixed fast and slow oscillatory phase are still visible, and are marked in panel C with the same color (dark gray, yellow and dark blue, respectively) as in Fig. 2C.

**Figure 11. [Figure S3].**
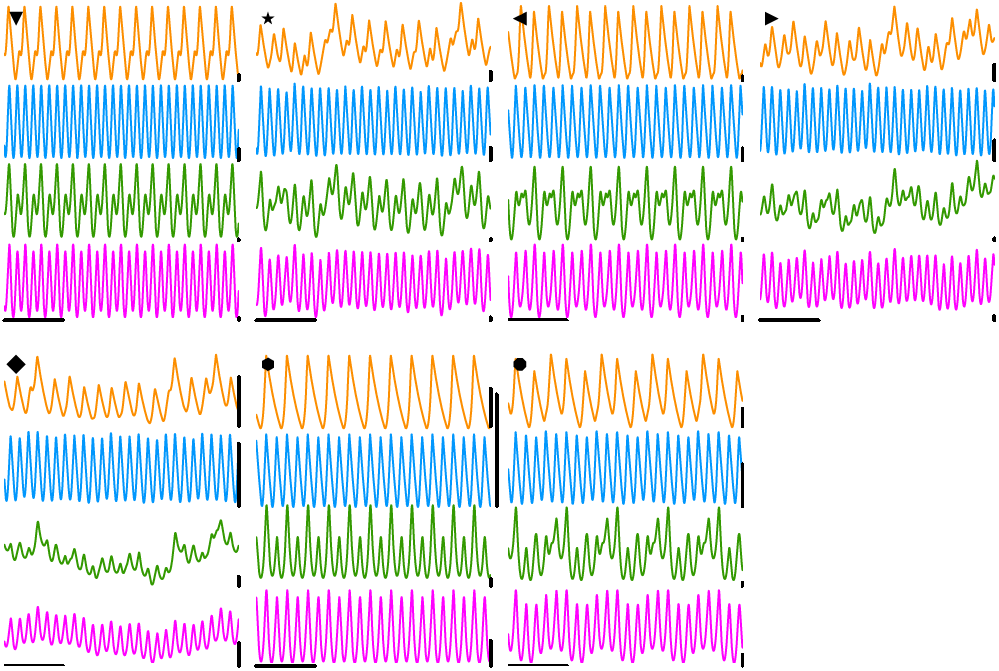
Additional example traces. Time-series of neural activity from the remaining working points marked in Fig. 2C. No contextual modulation is here applied (*I*^LGN^ = *I*^TD^ = 0, *I*^HOR^ =0)

**Figure 12 (following page). [Figure S4].**
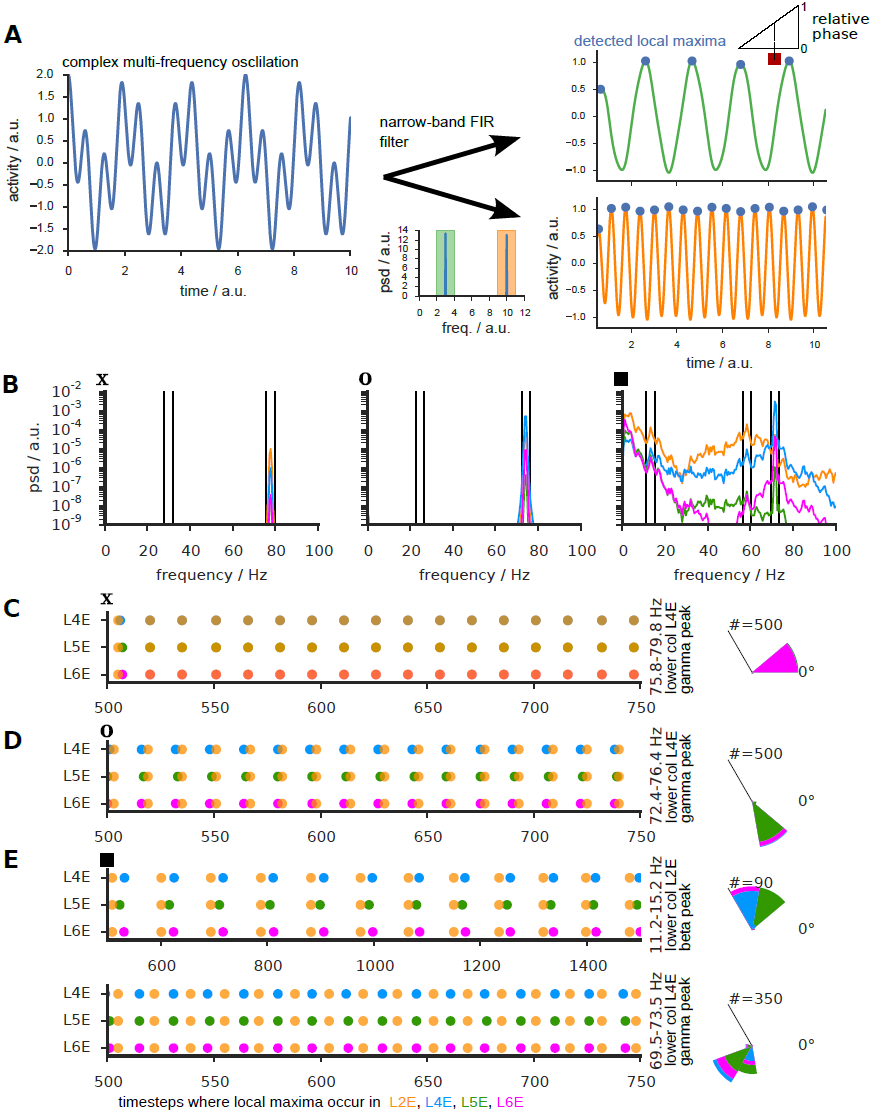
Definition of phase and phase-relations within a local circuit. (A) Illustration of our procedure for extracting frequency-resolved phase. After subsampling the original time series (for performance reasons) we apply a finite impulse response filter with a pass-band of ±2 Hz around a predominant frequency of the power spectral density, detect the local maxima in the filtered time series and interpolate linearly between the maxima to assign a relative phase to a target event, like a local maximum detected in the same way in another time series. **(B)** Power spectra for working points marked in Fig. 4 and limits of the narrow-frequency bands (fast and slow) used for filtering. **(C-E)** Time-stamps of local maxima in L23 (orange) relative to those in the other layers for a short time-segment. Different rows correspond to bla bla… ?. These time-stamps are extracted from the entire generated time-series and used to determine relative phases between oscillations in different layers and construct the histograms of their distribution in Figs 4B-D.

**Figure 13. [Figure S5].**
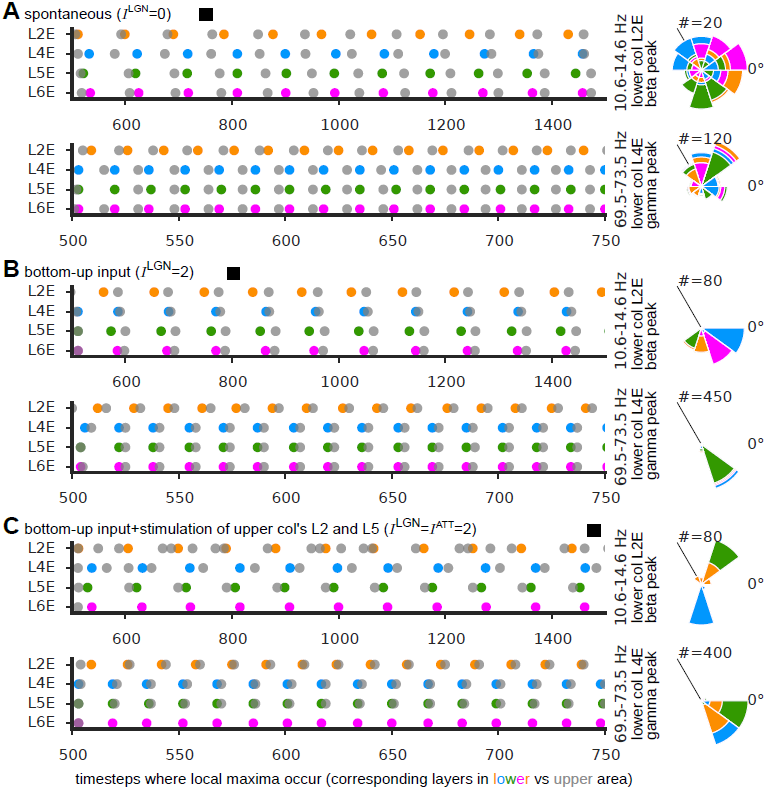
Phase-relations between interacting circuits at different stages of the cortical hierarchy. Time-stamps of local maxima in different layers of the lower canonic circuit (see color for legends). Different rows correspond to different layers. A grey dot correspond to time-stamps of the corresponding layer in the upper circuit.

**Figure 14 (following page). [Figure S6].**
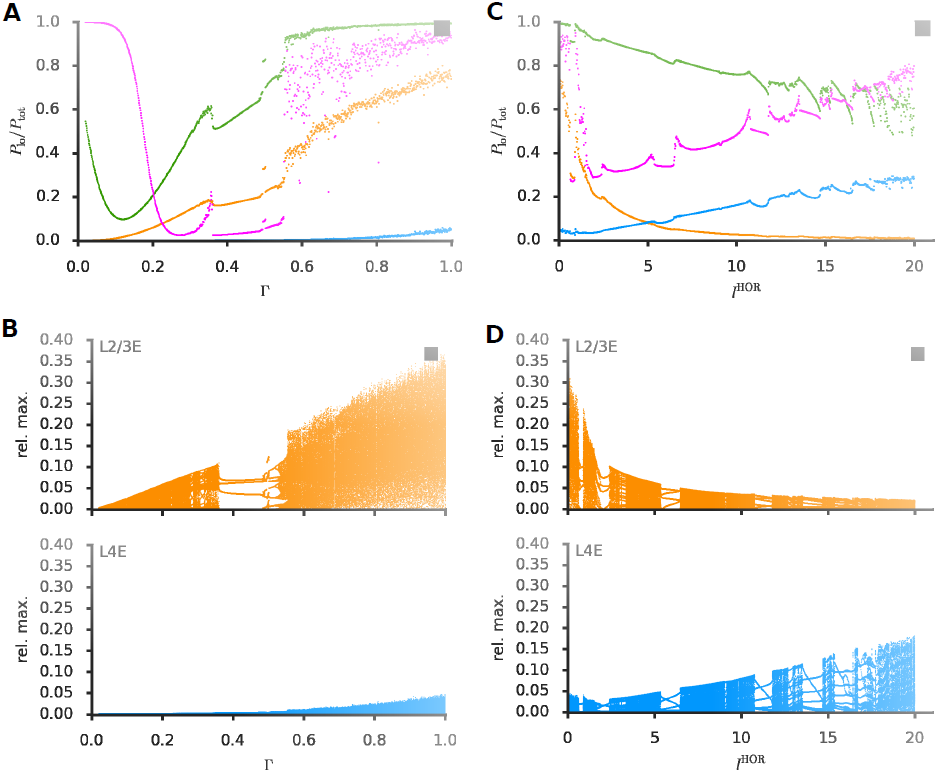
Chaotic entrainment between layers leads to broadband fast/slow oscillations. (A) Fractions of low power relative to total power for different layers in the ▪ working point (selected corresponding spectra are shown in Fig. 6A), as a function of the strength scale of inter-layer coupling Γ. **(B)** Corresponding bifurcation diagram, obtained plotting, for each value of Γ, a different dot for each observed value of oscillation maxima amplitudes. Cumulation of different dots for different Γ values results into tracing a tree structure, alternating sections with a discrete number of thin branches (corresponding to regular periodic oscillations, eventually period-doubled), with dense bands in which oscillation amplitudes cover continuous ranges of values (corresponding to chaotic oscillations). Details on the construction of these bifurcation diagrams can be found in [47, 48]. **(C–D)** Same as panels **A** and **B**, respectively, but as a function of the level of horizontal input to L23, *I*^HOR^. Discontinuities in the relative ratio of low power — associated to sharp qualitative transitions in the shape of power spectra (**A,C**)—, as well as the matching windows of regularity in bifurcation diagrams (**B,D**) support the idea that oscillations undergo a transition to chaos through a quasi-periodic route, as an effect of inter-layer entrainment. Note that the relative spread of amplitude values throughout the entrainment process is different for L4 and L23. For instance, in panel **B**, L4 oscillation amplitudes indeed display smaller relative modulations than L23’s oscillations. Note that in an ideal direct entrainment scenarion (rather than mutual) the bifurcation diagram for the driver would be a pure line, corresponding to the unique stable value of the driver’s fast oscillation amplitude. This different modulation amplitudes for L4 and L23 suggest that, for pure bottom-up drive in absence of modulation from horizontal inputs, L4 acts as an “effective driver” for L23. However, contextual modulation can change the effective role of these two layers in the inter-layer entrainment dynamics. Panel **D** shows indeed, that by increasing sufficiently *I*^HOR^, L23 can also be promoted to the role of effective driver, driving L4 into manifest chaos. Correspondingly, L4’s relative ratio of low power increases to higher values than for L23 when *I*^HOR^ is strong (panel **C**).

**Figure 15 (following page). [Figure S7].**
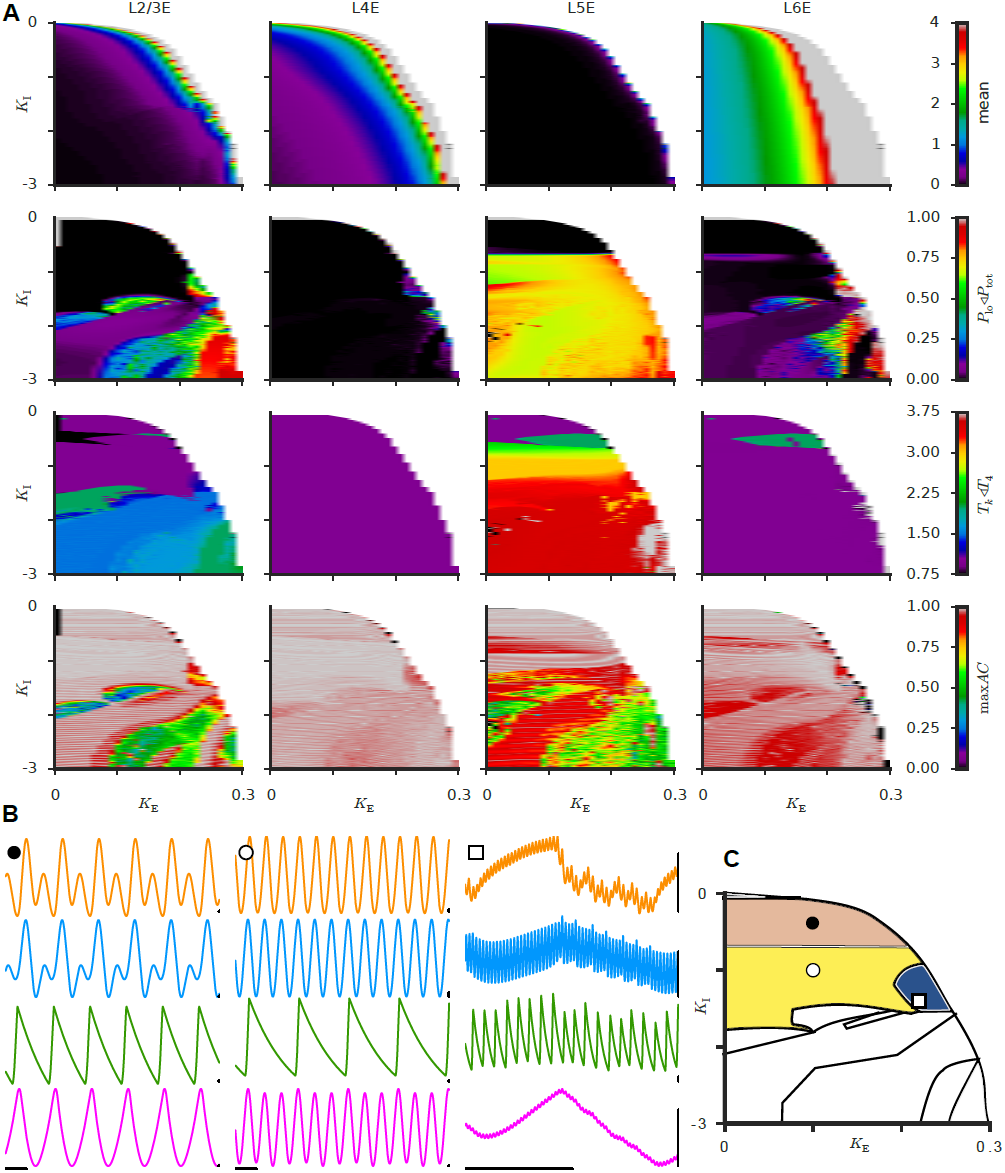
Dynamical repertoire for the alternative model with intrinsic slow frequency resonance. We present here dynamic profiles (panel **A**), time-series from selected working points (panel **B**) and a cartoon phase diagram (panel **C**), when the original model is modified to include an intrinsic resonance at slow frequency in L5. All panels are constructed analogously to Figs. 1 (*I*^LGN^ = 2, *I*^HOR^ = *I*^TD^ = 0). A fast/slow oscillatory regime originated by inter-layer entrainment (cf. Fig. 6B) is found at a similar position in the phase diagram as for the original model (denoted by a dark blue region, working point). Different types of periodic oscillatory regimes can be found, including one in which L4 and L23 oscillate at fast frequency, while L5 oscillates *endogenously* at a slow frequency close to its intrinsic resonance (◯ working point, “intrinsic fast/slow regime”) and one in which even L4 and L23 too develop a slow main period matching the one of L5, superposed with a smaller amplitude fast component (• working point, “all slow regime”)

**Figure 16 (following page). [Figure S8].**
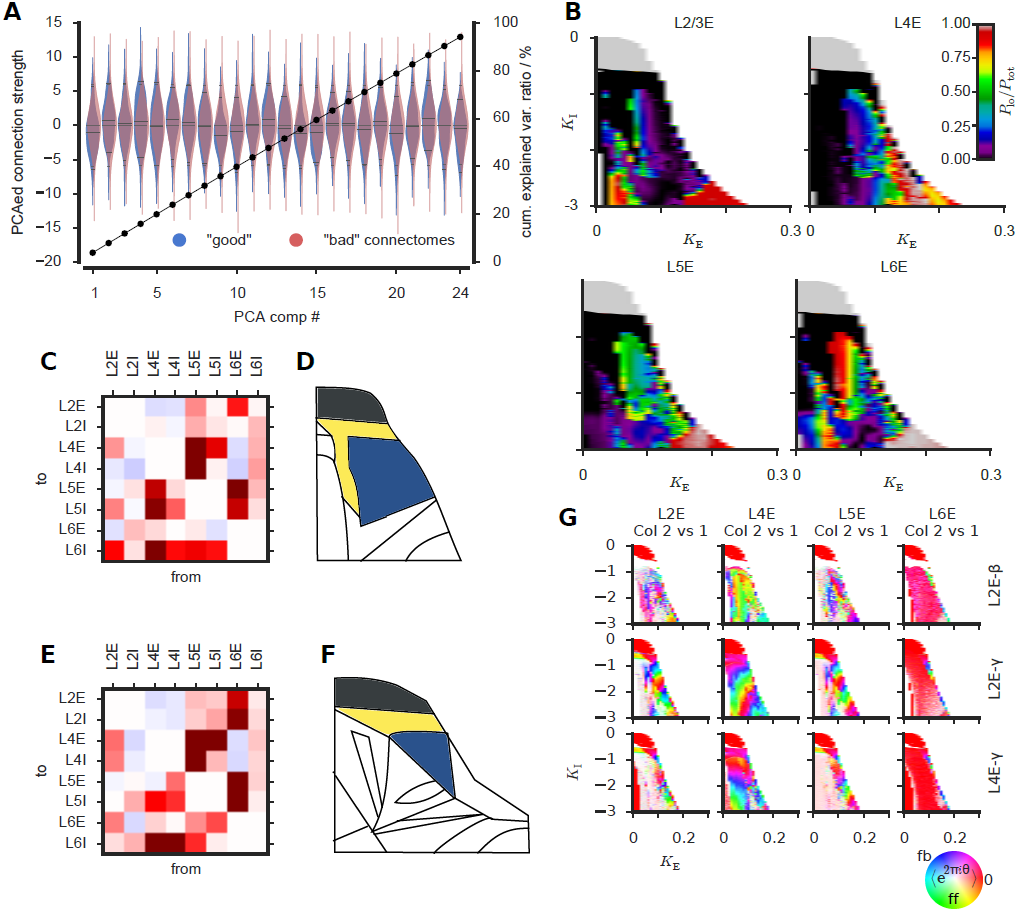
Additional results about randomized “good” connectomes. **(A)** Distributions of relevant PCA components for “good” and “bad” connectomes overlap. Violin widths (blue, “good” connectomes; red, other randomized connectomes) change along the left *y*-axis, describing distributions of the values of the first 24 PCA components (*x*-axis). Together, these 24 PCA components capture most of the variance, as indicated by the plot of cumulatively captured variance (line with black circles, right *y*-axis). Blue and red violins overlap almost completely indicating that PCA cannot separate “good” from “bad” connectomes. **(B)** Regime profiles for the fraction of low power for the “good” connectome illustrated in Figs. 8D,E. Panels **(C,D)** and **(E,F)**, show two more examples of randomized “good” connectomes analogous to Figs. 8D,E. **(G)** We coupled two local circuits with the alternative “good” connectome of Figs. 8D,E, to mimic interaction between cortical circuits at different hierarchical levels, as in Fig. 1F and 5. Shown here are the dynamic profiles for the relative phase between L23 of the lower circuit and L23 of the upper circuit in presence of bottom-up input only. Although we did not attempt a systematic study of the effects of contextual modulation or attention, these profiles reveal a richness of possible frequency-dependent locking patterns comparable with Fig. 5D.

